# Insect-specific viruses regulate vector competence in *Aedes aegypti* mosquitoes via expression of histone H4

**DOI:** 10.1101/2021.06.05.447047

**Authors:** Roenick P. Olmo, Yaovi M. H. Todjro, Eric R. G. R. Aguiar, João Paulo P. de Almeida, Juliana N. Armache, Isaque J. S. de Faria, Flávia V. Ferreira, Ana Teresa S. Silva, Kátia P. R. de Souza, Ana Paula P. Vilela, Cheong H. Tan, Mawlouth Diallo, Alioune Gaye, Christophe Paupy, Judicaël Obame-Nkoghe, Tessa M. Visser, Constantianus J. M. Koenraadt, Merril A. Wongsokarijo, Ana Luiza C. Cruz, Mariliza T. Prieto, Maisa C. P. Parra, Maurício L. Nogueira, Vivian Avelino-Silva, Renato N. Mota, Magno A. Z. Borges, Betânia P. Drumond, Erna G. Kroon, Luigi Sedda, Eric Marois, Jean-Luc Imler, João T. Marques

**Author notes:** These authors contributed equally to this work (Roenick P. Olmo, Yaovi M. H. Todjro, Eric R. G. R. Aguiar).

## Abstract

*Aedes aegypti* and *Aedes albopictus* are major mosquito vectors for arthropod-borne viruses (arboviruses) such as dengue (DENV) and Zika (ZIKV) viruses. Mosquitoes also carry insect-specific viruses (ISVs) that may affect the transmission of arboviruses. Here, we analyzed the global virome in urban *Aedes* mosquitoes and observed that two insect-specific viruses, Phasi Charoen-like virus (PCLV) and Humaita Tubiacanga virus (HTV), were the most prevalent in *A. aegypti* worldwide except for African cities, where transmission of arboviruses is low. Spatiotemporal analysis revealed that presence of HTV and PCLV led to a 200% increase in the chances of having DENV in wild mosquitoes. In the laboratory, we showed that HTV and PCLV prevented downregulation of histone H4, a previously unrecognized proviral host factor, and rendered mosquitoes more susceptible to DENV and ZIKV. Altogether, our data reveals a molecular basis for the regulation of *A. aegypti* vector competence by highly prevalent ISVs that may impact how we analyze the risk of arbovirus outbreaks.

## Main Text

Arthropod-borne viruses (arboviruses) transmitted by mosquitoes, such as DENV, ZIKV and chikungunya (CHIKV) viruses, are a great threat to human health worldwide (*1*). DENV alone is estimated to cause 400 million infections per year, leading to approximately 20,000 deaths (*2*). DENV transmission has increased several folds over the past decades and newly emerged mosquito borne viruses such as CHIKV and ZIKV have become important (*1, 3*). Increased transmission of arboviruses has been enabled by the global spread of *Aedes aegypti* and *Aedes albopictus* that are their major urban mosquito vectors. Globalization, urbanization and climate change have positively impacted the distribution of these *Aedes* mosquitoes worldwide (*4–6*). Importantly, we still lack vaccines or treatments for most arboviral diseases (*1*).

Arboviruses circulate primarily in enzootic cycles involving sylvatic mosquitoes and vertebrate animals causing rare accidental human infections (*6, 7*). However, arboviruses may become able to infect humans efficiently and be maintained by human-mosquito-human transmission in nature that will no longer require animal reservoirs (*1*). In this scenario, virologic surveillance of urban *Aedes* mosquitoes can lead to early identification of circulating viruses and help raise preparedness to prevent outbreaks (*8*). However, studies monitoring the collection of viruses, the virome, in *Aedes* mosquitoes have mostly identified a large diversity of insect-specific viruses (ISVs) (*9–13*). ISVs do not infect vertebrates but can affect the capacity of the mosquito to be infected, maintain and transmit arboviruses, usually referred to as vector competence (*8, 14, 15*). However, most data on ISV-arbovirus interactions have been obtained in cell lines using closely related viruses that may compete for common resources (*16–26*). In addition, effects were often reported for ISVs that belong to the same family as arboviruses and are likely to compete for common resources, a phenomenon referred to as superinfection exclusion (*17, 19, 23*). Mechanistic understanding of how ISVs affect arboviruses beyond the competition between related viruses remains scarce, especially *in vivo*, and we still lack evidence for interactions in natural conditions. In this regard, the presence of ISVs could help explain differences in vector competence between mosquito populations thus contributing to outbreaks.

Here, we characterize the virome of urban *Aedes* mosquitoes worldwide and provide novel mechanistic understanding of the interactions between ISVs and arboviruses in natural conditions.

## Results

*A. aegypti* and *A. albopictus* mosquitoes are the most prolific and widespread vectors for arboviruses (*5, 8*). To assess the collection of viruses found in these mosquitoes, we took advantage of a network of field researchers wordlwide. Adult *A. aegypti* and *A. albopictus* mosquitoes were collected directly from the field in 12 different sites from 6 countries on 4 continents (**Figure 1a**). In total, 815 adult mosquitoes were pooled according to species, location and date of collection resulting in 91 samples derived from 69 *A. aegypti* and 22 *A. albopictus* pools or individual insects (**Supplementary Table 1**). These 91 samples were used to construct small RNA libraries that were sequenced and analyzed using a metagenomic strategy, previously described by our group (*9*), optimized to detect viruses (**Figure 1b**). In total, we were able to assemble 7260 contiguous sequences (contigs) larger than 200 nt from the 91 individual libraries (**Figure 1b**). A summary of the assembly results is shown in **Supplementary Table 2**. Out of these 7260 contigs assembled, 1448 contigs were identified as putative viral sequences by sequence similarity searches against non-redundant nucleotide and protein databases (NT and NR, respectively) at GenBank (**Figure 1b**, **Supplementary Table 3**). Although the number of contigs assembled per library varied greatly, we observed high abundance and diversity of viral contigs in most samples (**Figure 1c**). Comparing results from the two mosquito species, the percentage of viral contigs was strikingly smaller in *A. albopictus* libraries compared to *A. aegypti* (**Figure 1c**). In addition, we noted more variation in the number of assembled contigs and larger proportions of unknown contigs in libraries from *A. albopictus*, probably because this species is less studied compared to *A. aegypti* (**Figure 1c**).

**Fig. 1.**
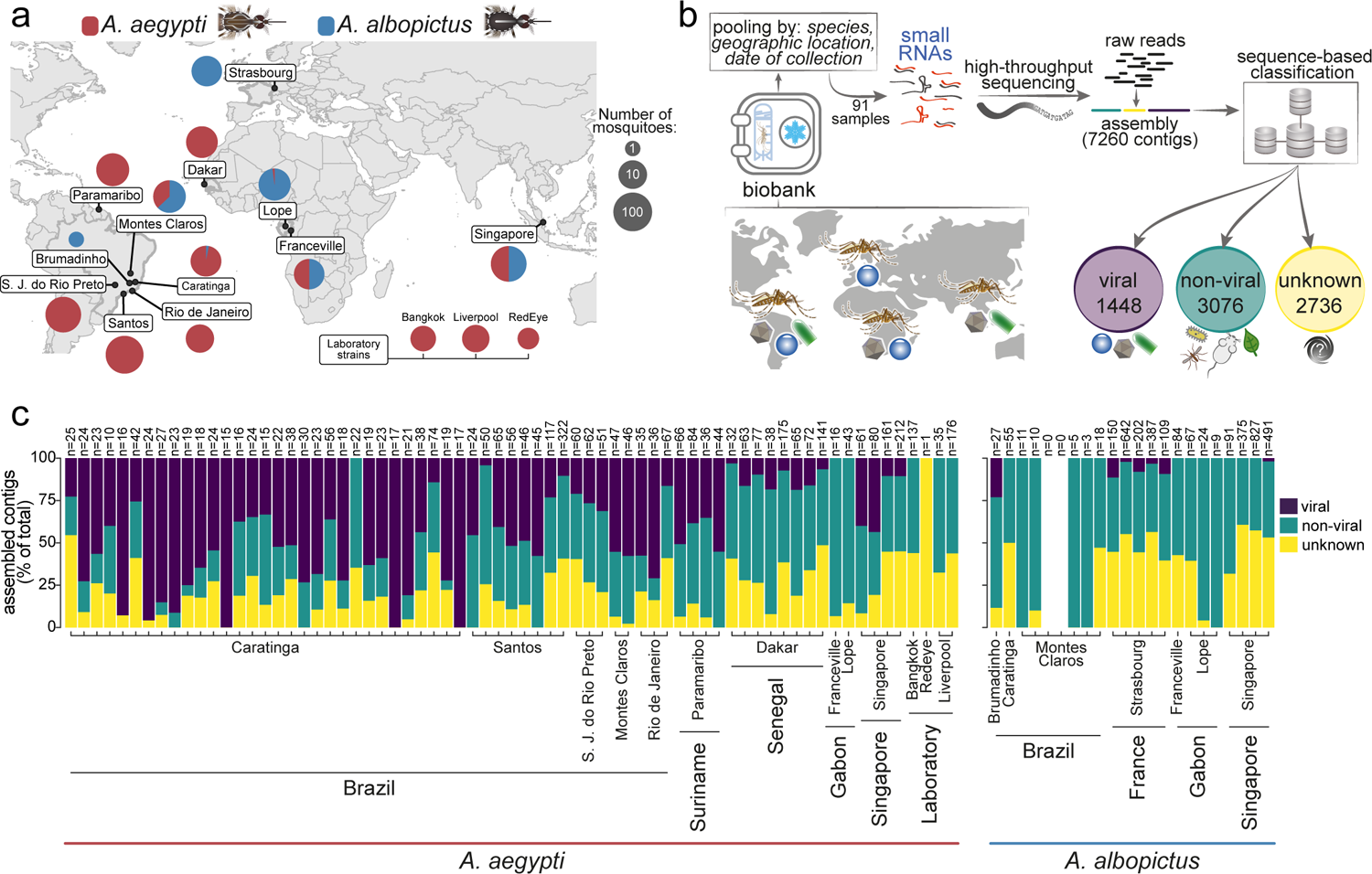
Analysis of the global virome of *A. aegypti* and *A. albopictus* mosquitoes. **a**, World map indicating sites of mosquito collection. *A. aegypti* and *A. albopictus* are shown in red and blue, respectively. Pie charts show the proportion of species at each collection site. Adult mosquitoes were captured either using traps or human baits. Laboratory strains of mosquitoes analyzed in this work are indicated at the bottom. **b**, Overview of our analysis pipeline. Captured mosquitoes were morphologically identified by species and stored in a biobank of RNA samples. Samples were pooled and used to prepare small RNA libraries for high-throughput sequencing that were analyzed using our metagenomic pipeline. Assembled contigs were classified into viral and non-viral sequences based on similarity against reference databases. Sequences without any similarity to known references in the database are indicated as unknown. **c**, Individual results from our metagenomic analysis for each of the 91 small RNA libraries in this study. The total number of contigs and the proportion of viral, non-viral and unknown contigs are indicated.

Most animal genomes contain integrated viral sequences known as endogenous viral elements (EVEs) that are transcribed and generate small RNAs (*27–29*). In order to discriminate sequences derived from *bona fide* viruses and EVEs, we took advantage of the small RNA profile associated with ORF analysis and contig size to discriminate sequences derived from exogenous viruses from EVEs (**Figure 2a**) (*30*). This filter identified 446 putative EVE sequences that were removed from the initial 1448 viral contigs (**Figure 2a**). The remaining 1002 viral contigs representing putative viruses were grouped into 158 unique clusters based on sequence similarity to remove redundancy (**Supplementary Table 4**). In 19 of the 158 clusters, contigs had sections that showed significant similarity to two viruses and had distinct patterns of sequence coverage. Our results suggested that these were misassemblies and 19 contigs were removed from further analyses (**Figure 2a**). Contigs representing the remaining 139 clusters were used to evaluate co-occurrence in the 91 independent small RNA libraries from *A. aegypti* and *A. albopictus* (**Supplementary Figure 1**). The occurrence of contigs in each library was indicated by the normalized number of small RNA reads mapped to each reference. Across the 91 small RNA libraries, contigs that consistently co-occurred and shared similar expression profiles were considered probable fragments from the same viral genome (**Figure 2a**). This analysis yielded a total of 12 clusters of co-occurring contigs and 3 single contigs that had no additional partners. Contigs from each of the 12 clusters were further analyzed based on the closest reference sequence to determine the putative organization of fragments along the viral genome. This analysis showed that clusters #2 and #3 contained contigs belonging to the same virus, PCLV, and were considered together for further analyses (**Supplementary Figure 1**). In order to classify the 11 clusters and 3 single contigs, we focused on the ones that represented sequences encoding viral polymerases. We were able to identify clear polymerase sequences in each of the 11 clusters and in one out the 3 single contigs. These contigs with similarity to viral polymerases were compared to the closest sequence in GenBank suggesting that they represented at least 12 different viruses (**Figure 2b**). Seven viruses were previously known and had all been previously identified as ISVs. Out of these 7, three remain unclassified while the other 4 belong to the *Phenuiviridae*, *Xinmoviridae*, *Bunyaviridae* and *Flaviviridae* families (**Figure 2b**). No known arboviruses were detected in our metagenomic analysis. The five remaining viral polymerase sequences showed similarity below 95% to the closest known reference, which suggested they could represent new viral species. Phylogenetic analyses confirmed that they are likely new viruses (**Supplementary Figure 2**), belonging to the *Partitiviridae*, *Totiviridae*, *Rhabdoviridae*, *Narnaviridae* and *Virgaridae* families (**Figure 2b**). These new viruses were named according to their classification (**Figure 2b**). All 5 new viruses were most closely related to known ISVs (**Supplementary Figure 2**), but their final classification requires biological characterization.

**Fig. 2.**
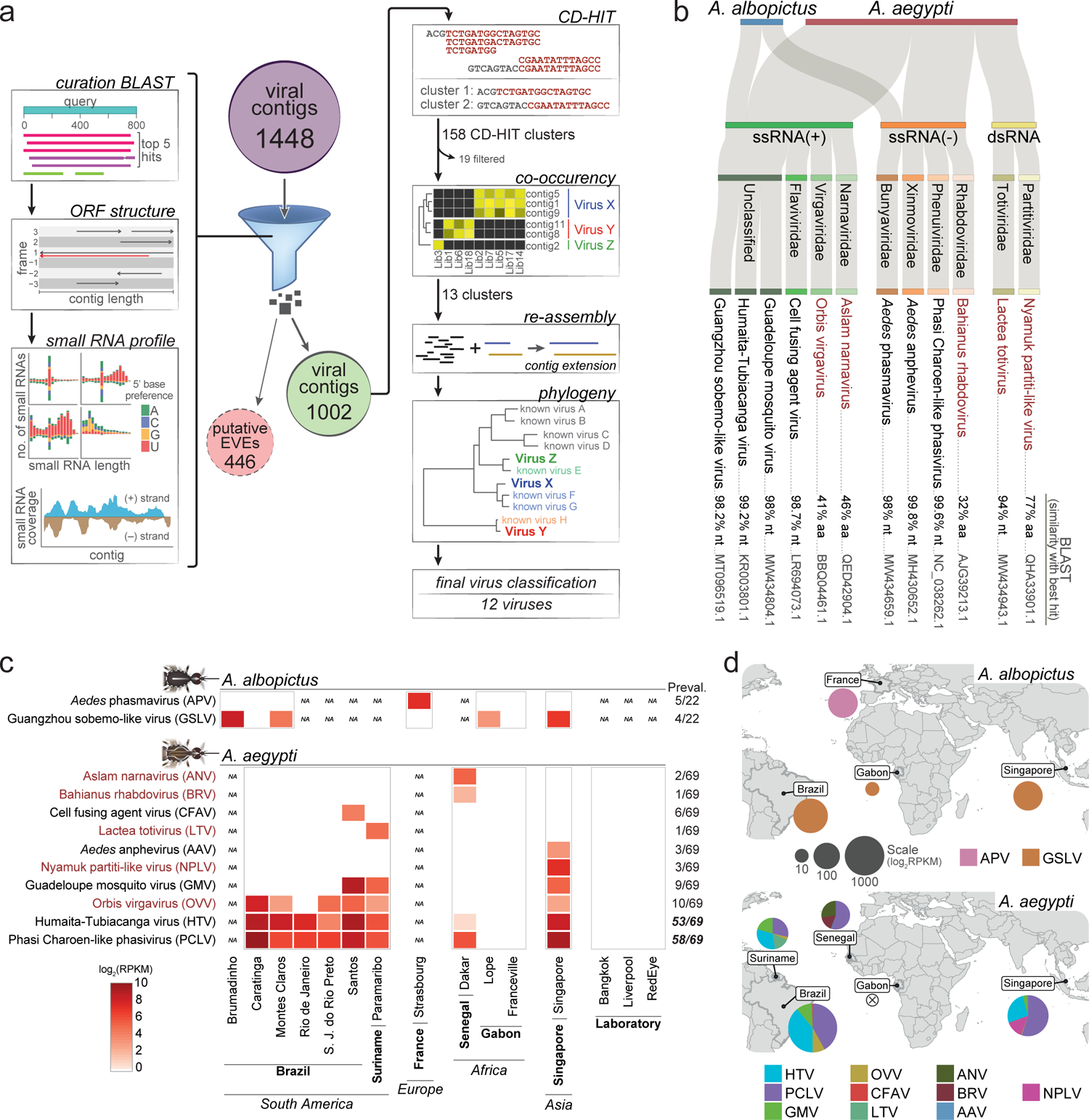
A highly diverse and distinct virome in *A. aegypti* and *A. albopictus*. **a**, Overview of the strategy for manual curation of viral contigs to confirm the origin and filter sequences potentially derived from EVEs. Curation consisted of BLAST search for similar viral sequences, inspection of ORF structure including continuity and overall extension throughout the contig, and the evaluation of the small RNA profile including symmetrical accumulation of RNAs with 20-23 nt length that mapped to each contig strand with no 5’ base preference (siRNA signature) or accumulation of 24-29 nt RNAs with 5’ U preference (piRNA signature); and overall contig coverage by small RNAs. Remaining viral contigs are clustered by sequence similarity and co-ocurrence in each library to identify groups of contigs that belong to the same virus. Re-assembly was perfomed within these groups and resulting contigs were analyzed for the presence of domains. Potential polymerases were identified and used to classify viruses based on sequence similarity and phylogeny. **b**, Host, virus genomic organization, family, and closest reference on GenBank identified by BLAST similarity searches for each of the 12 viruses identified in our datasets. New viral species are indicated in red while previously known viruses are in black. Sequence similarity and accession number according to the closest viral sequence at the nucleotide (nt) or protein (aa) levels are indicated. **c**, Viral load shown as a heatmap for each of the twelve viruses in mosquito populations from each collection site or laboratory strains. In the heatmap, white indicates absence of the virus in the sample and NA the absence of the sample. Prevalence of each virus is shown on the right as number of samples with detectable virus over the total. Number of individuals per pool and number of species per collection site are described in detail on the Supplementary Table 1. **d**, Pie charts representing the overall burden of virus and viral diversity for *A. aegypti* and *A. albopictus* populations indicated in the world map. The icon “absent” indicates that no viral contigs have been identified in a species at a specific site.

All 12 identified viruses, 7 known and 5 new, had RNA genomes, either single-stranded (of positive and negative polarity) or double-stranded (**Figure 2b**). The small RNA profile observed for the 12 viruses shows clear production of siRNAs (**Supplementary Figure 3**), which results from the activity of the RNAi pathway during active viral replication in the mosquito host (*9, 30–34*). Viruses detected in *Aedes* mosquitoes were strictly species-specific and often associated with specific locations (**Figure 2c**). Out of the 12 identified viruses, 10 were found in *A. aegypti* and 2 in *A. albopictus* suggesting a less diverse virome in the latter even accounting to a lower number of samples. Indeed, looking at the diversity of the mosquito virome per country, a single virus species was detected in each *A. albopictus* population while 4-6 different viruses were present in *A. aegypti* (**Figure 2d**). In addition, comparing different mosquito species that were collected from the same site in Caratinga, Montes Claros, Lope, Franceville and Singapore, we observe that *A. aegypti* had higher virome diversity than *A. albopictus* in 4 out of 5 cases (**Figure 2c**).

Analyzing the number of small RNAs mapping to each virus contig as a proxy for abundance, we observed that viral loads varied for the same virus in different locations and also between different viruses (**Supplementary Figure 4**). Three locations had 5 or more viruses circulating in the local *A. aegypti* population: Santos, Paramaribo and Singapore (**Figure 2c**). Notably, these are all port cities that might have a continuous influx of mosquitoes. On the other hand, three different strains of laboratory *A. aegypti* mosquitoes carried no viruses in our analysis (**Figure 2c**). Most viruses that we identified, 7 in total, were found at single sites but 5 were present in at least 2 continents (**Figure 2d**). No viruses were found with a prevalence higher than 20% in *A. albopictus* samples and only one was present in multiple sites. In *A. aegypti*, 8 out of the 10 viruses were also found at low prevalence, less than 20% of the samples although two known ISVs, PCLV and HTV, were present in over half of the samples (**Figure 2c**). HTV and PCLV also had the highest viral loads of all viruses we found in either mosquito species (**Supplementary Figure 4**). These viruses were only absent in *A. aegypti* collected in two locations, Lope and Franceville, both in Gabon, Africa (**Figure 2c**). Other *A. aegypti* samples from Africa (Dakar, Senegal) had the lowest overall viral loads for PCLV and HTV (**Supplementary Figure 4**). Notably, transmission of arboviruses such as DENV and ZIKV is also low in African countries (*1, 2*). In contrast, high prevalence of HTV and PCLV was observed in areas with high transmission of arboviruses in Asia and South America (*1, 2*).

In order to examine the spatiotemporal dynamics of the virome in wild mosquitoes, we choose to focus on one of the 12 sites used for the metagenomic analysis. Here, we took advantage of a biobank of individual mosquitoes, 515 *A. aegypti* and 24 *A. albopictus*, collected for an entire year in the city of Caratinga in the southeast of Brazil (**Figure 3a**). This dataset was previously used to assess DENV circulation in an endemic urban area (*35*). In our metagenomic analysis, we detected 3 viruses in mosquitoes from Caratinga: Orbis virgavirus (OVV), HTV and PCLV (**Figure 2c**). By analyzing individual mosquitoes, we confirmed our metagenomic results using RT-qPCR; OVV, HTV and PCLV were only detected in wild *A. aegypti* and not in *A. albopictus,* even though these two species were captured often in the same traps (**Figure 3a**). OVV was only detected in 3 individuals out of 515 tested while HTV and PCLV were found at 61% and 83% prevalence in the mosquito population, respectively (**Figure 3b,c**). This analysis confirmed that HTV and PCLV but not OVV were found at high prevalence in natural populations. Due to the small number of mosquitoes with OVV, this virus was not included in further analyses. HTV and PCLV were found at similar high prevalence throughout the year and independently of the location of collection within the city (**Figure 3d** and **Supplementary Figure 5**). Interestingly, we observed a strong positive association between the presence of HTV and PCLV in mosquitoes (chi-squared test *p*-value = 2.28e-16), suggesting the co-infection might be advantageous for each virus.

**Fig. 3.**
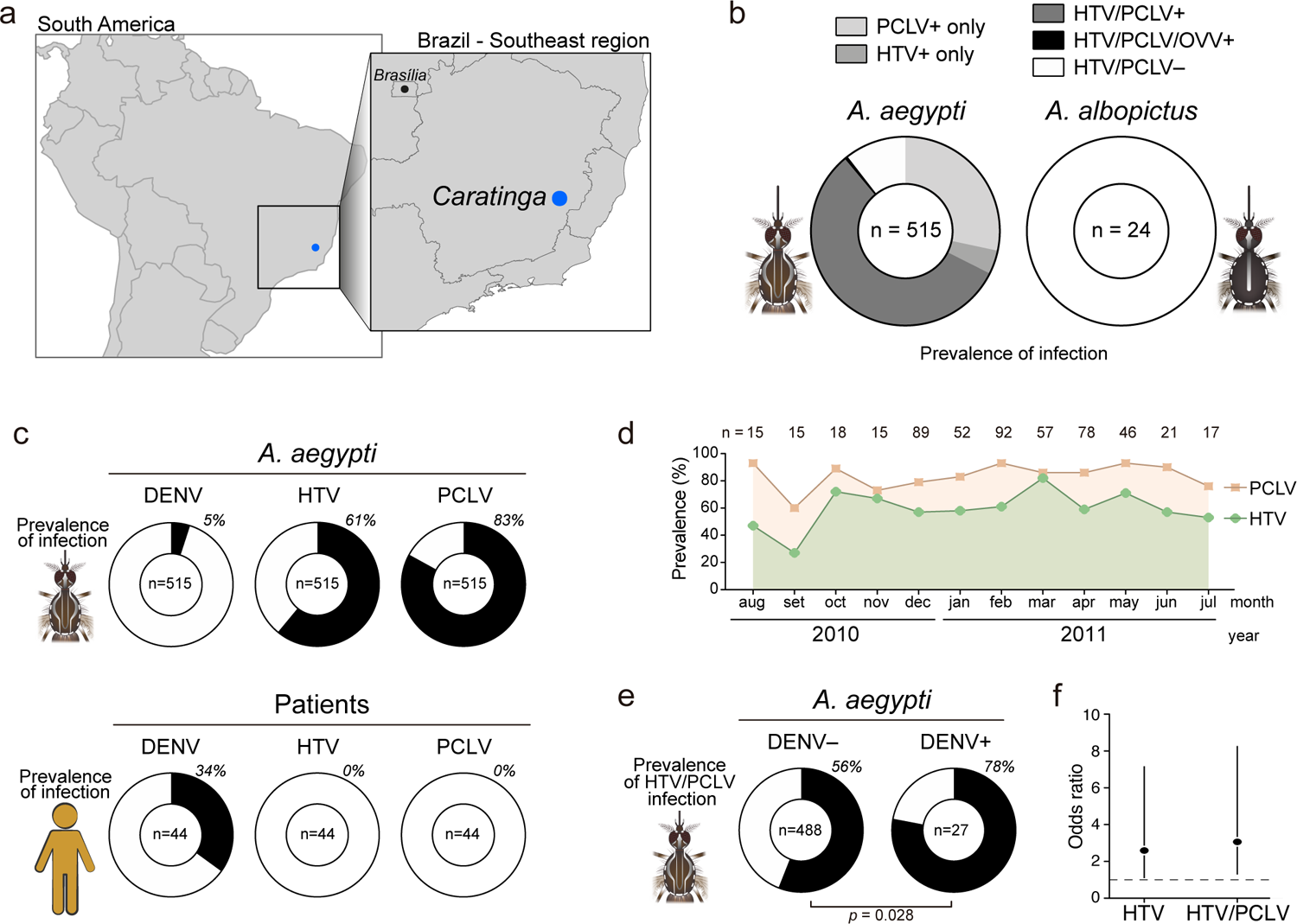
Spatiotemporal dynamics of the mosquito virome in a wild mosquito population shows positive interactions between HTV and PCLV with DENV. **a**, Location of the study site, the city of Caratinga in the Southeast of Brazil. **b**, Prevalence of individual and co-infections by OVV, HTV and PCLV tested by RT-qPCR. **c**, Prevalence of DENV, HTV and PCLV separately in individual *A. aegypti* mosquitoes and human blood samples accessed by RT-qPCR. **d**, Monthly prevalence of HTV and PCLV separately in individual *A. aegypti* mosquitoes. **e**, Prevalence of HTV and PCLV co-infection in DENV infected (DENV+) and DENV non-infected (DENV-) mosquitoes. Statistical significance was determined by two-tailed Fisher’s exact test. **f**, Likelihood of DENV infection in mosquitoes carrying HTV or PCLV and HTV shown by odds ratio.

HTV and PCLV are presumed to be ISVs although data on their biological characterization remain scarce (*9, 20, 22*). To assess the possibility that HTV and PCLV could be transmitted to humans, we analyzed the presence of these viruses in human blood samples collected concomitantly with mosquitoes. These blood samples were previously analyzed to correlate DENV circulation in mosquitoes and humans in Caratinga (*35*). In this comparison, DENV was detected in less than 5% of mosquitoes but over 30% of human patients (**Figure 3c**). In contrast, HTV or PCLV were not detected in human blood samples despite their high prevalence in mosquitoes at the same time, suggesting they are not able to productively infect humans (**Figure 3c**). HTV and PCLV were also not able to grow in mammalian cell lines such as Vero cells (**Supplementary Figure 6**). Together these results reinforce the idea that HTV and PCLV are ISVs.

Although ISVs are not infectious to humans, they have been shown to affect how arboviruses are transmitted. In our dataset, we observed a strong enrichment of HTV and PCLV in mosquitoes carrying DENV (**Figure 3E**). In order to look further into this association, we applied a zero-inflated binomial model that returned two instances with significantly different covariates with DENV in mosquitoes: HTV alone and the co-infection by PCLV and HTV. In this model, PCLV alone had no statistically significant increase in the odds of DENV presence. The odds ratios (OR) of the models show that the presence of PCLV and HTV is associated with approximately 200% increase in the odds of having mosquitoes with DENV (OR 3.06, 95% Confidence Interval 1.29-8.46) while HTV alone also had a significant but weaker association (OR 2.59, 95% CI 1.09-7.16) (**Figure 3f**). However, due to strong association between HTV and PCLV in mosquitoes, it is hard to dissect the contribution of each virus. Thus, our observations using field samples strongly suggest a positive interaction between two ISVs, HTV and PCLV, with an arbovirus, DENV.

To further analyze the positive association between HTV and PCLV with DENV infection observed in Caratinga, we collected wild mosquitoes and reared them in the laboratory. From the original wild population, we used individual females to derive mosquito lines that were either free of any virus or carried both HTV and PCLV (see methods for details). We were unable to isolate lines carrying just HTV or PCLV, reinforcing the strong association we observed between these mosquitoes in samples directly collected in the wild. Considering this association, we carried out our functional analysis between mosquito lines that were virus free or co-infected by HTV and PCLV. When given an infectious blood meal in the laboratory, mosquitoes carrying HTV and PCLV had similar DENV prevalence and viral loads in the midgut at 4, 8 and 14 days post feeding compared to virus free individuals (**Figure 4a,b**). At 14 days post feeding, we observed a trend of higher viral loads and prevalence of DENV in the midgut of mosquitoes carrying HTV and PCLV but that was not statistically significant (**Figure 4b**). At this time point, the midgut probably reflects the status of systemic infection rather than the initial viral replication in this tissue. Indeed, analyzing carcass infection as a proxy of systemic dissemination, we observed a significant increase in DENV levels at 8 and 14 days post feeding in mosquitoes carrying HTV and PCLV compared to virus free controls (**Figure 4c**). A similar pattern of increased susceptibility in mosquitoes with HTV and PCLV was observed for ZIKV (**Figure 4d-f**). However, in the case of ZIKV, we observed a significant increase in viral RNA levels at 4, 8 and 14 days post feeding in the midgut of mosquitoes carrying HTV and PCLV compared to virus free controls (**Figure 4e**). In the carcass, we observed earlier dissemination and significantly higher viral loads at 8 days post feeding in the presence of HTV and PCLV. Overall, the kinetics and rates of infection for ZIKV were higher compared to DENV and could explain the different effect of HTV and PCLV. Nevertheless, lines derived from wild mosquitoes with HTV and PCLV show increased susceptibility to DENV and ZIKV. The selection of wild mosquitoes based on the presence of HTV and PCLV could result in lines with different genetic backgrounds. Thus, rather than the cause of the phenotype, the presence of ISVs could be associated with a genetic background more susceptible to DENV and ZIKV. In order to verify whether HTV and PCLV have a direct effect on the susceptibility to arboviruses, we artificially infected laboratory mosquitoes with HTV and PCLV (**Figure 4g**). Notably, HTV and PCLV loads and tissue tropism during artificial injection of laboratory mosquitoes were similar to naturally infected individuals after 8 days post injection (**Supplementary Figure 7**). Artificially infected laboratory mosquitoes had increased systemic ZIKV RNA levels at 8 days post feeding compared to controls, similar to what we observed for lines carrying HTV and PCLV derived from wild populations (**Figure 4g-i**). These results suggest that DENV and ZIKV have consistently higher systemic infection rates in mosquitoes carrying HTV and PCLV compared to virus-free controls. In order to look further into the effect of HTV and PCLV during the systemic phase of infection, we directly injected ZIKV into the mosquito hemocoele, which bypasses the stage of midgut infection (**Supplementary Figure 8**). Mosquito lines derived from wild populations carrying HTV and PCLV injected with ZIKV had increased viral replication in the carcass compared to virus-free controls (**Supplementary Figure 8a-c**). The same was observed when laboratory mosquitoes were artificially infected with HTV and PCLV before being injected with ZIKV (**Supplementary Figure 8d-f**). These results show a direct impact of HTV and PCLV infection on the susceptibility of mosquitoes to systemic infection with DENV and ZIKV.

**Fig. 4.**
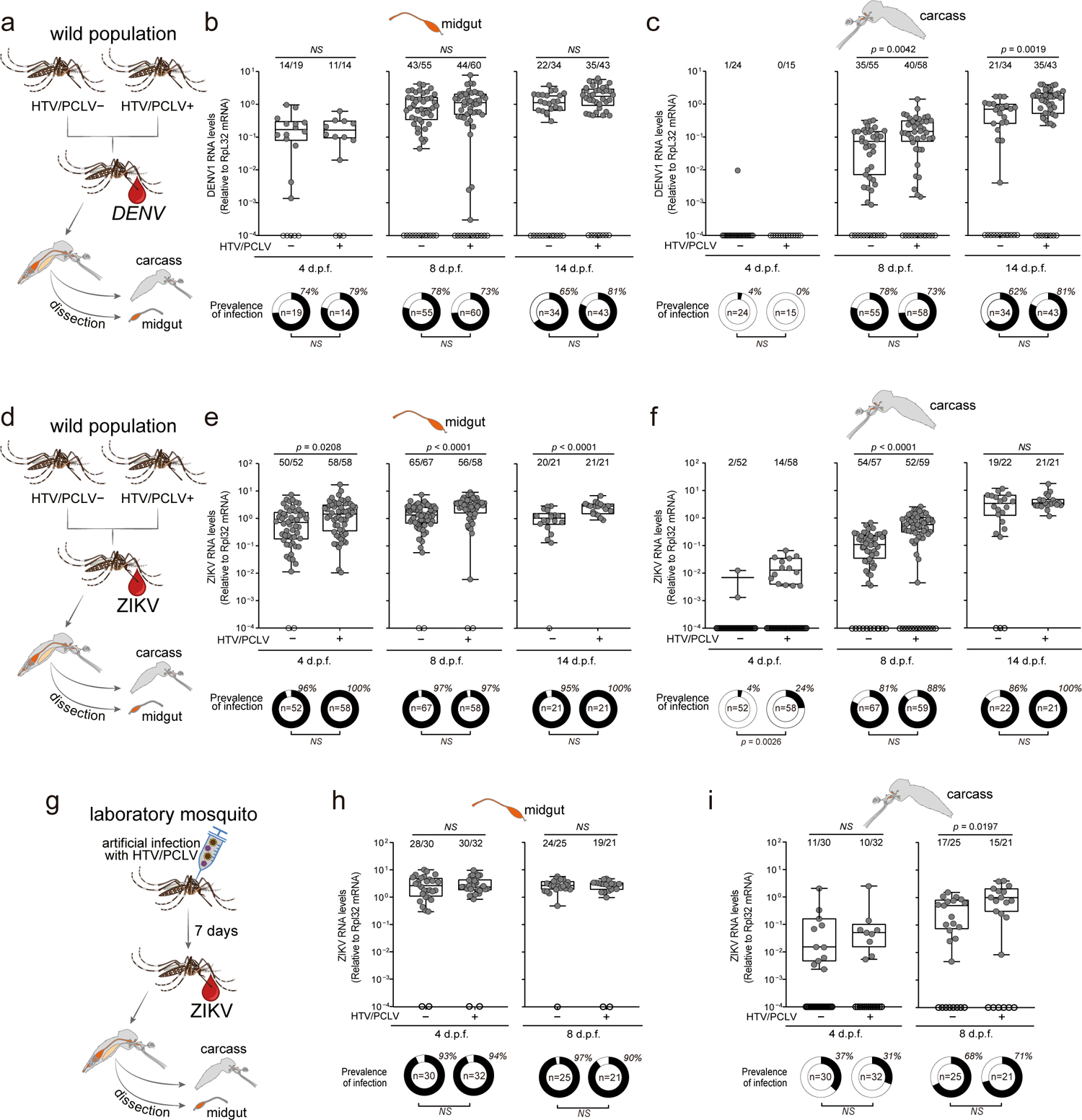
HTV and PCLV facilitate DENV infection in *A. aegypti* mosquitoes. **a-f**, Strategy to evaluate the interference of HTV and PCLV on DENV (**a-c**) or ZIKV (**d-f**) infection and replication in natural populations of mosquitoes. HTV/PCLV infected or virus free mosquito populations were exposed to DENV (**a**) or ZIKV (**d**) on a blood meal. Viral loads and prevalence of infection were measured in the (**b,e**) midgut and (**c,f**) carcass of mosquitoes at 4, 8 and 14 days post feeding. The prevalence of infection in each group is shown below the plots. **g-i**, Laboratory mosquitoes were infected artificially with HTV and PCLV and 7 days later were given a blood meal containing ZIKV (**g**). Viral loads and prevalence of infection were measured in the (**h**) midgut and (**i**) carcass of mosquitoes at 4 and 8 days post feeding. The prevalence of infection in each group is shown below plots. *d.p.f.* – days post feeding, *NS* – non-significant. In ***b***, ***c***, ***e***, ***f***, ***h***, and ***i***, error bars represent median, maximum, minimum, and interquartile range of samples above the threshold for virus detection. Statistical significance was determined by two-tailed Mann–Whitney U-test. Numbers of infected midguts (***b***,***e***,***h***) or carcasses (***c***,***f***,***i***) over the total number tested are indicated above each column. Each dot represents an individual sample. Statistical significance of prevalence of infection was determined by two-tailed Fisher’s exact test.

In order to shed light into the biological mechanisms by which ISVs affect the susceptibility of *A. aegypti* mosquitoes to systemic dissemination of DENV and ZIKV, we analyzed the transcriptome of mosquitoes. The transcriptome of DENV infected and non-infected individuals from groups of mosquitoes carrying HTV and PCLV or virus-free controls were compared using Gene Set Enrichment analysis (GSEA) (*36*) (**Figure 5a**). We analyzed the carcass of mosquitoes infected by DENV at 8 and 14 days post feeding when the presence of HTV and PCLV had the strongest effect (**Figure 5a**). This analysis identified 7 biological processes that were significantly affected both by the presence of HTV and PCLV and DENV infection in at least one time point (**Figure 5b**). Notably, in all these cases, the 7 processes were downregulated during DENV infection and upregulated by the presence of HTV and PCLV as indicated by the enrichment score (NES) (**Figure 5b**). Out of these 7 processes, 4 were significantly affected in at least 6 out of 8 comparisons: *nucleosome*, *nucleosome assembly*, *DNA templated transcription initiation* and *protein heterodimerization activity* (**Figure 5b**). Analysis of genes responsible for the significant enrichment of these 4 biological processes showed that they were overwhelmingly shared (**Figure 5c**). Furthermore, histones represented the majority of genes differentially regulated by DENV infection and the presence of HTV and PCLV with histone H4 being enriched in most conditions (**Figure 5c**). Thus, we used RT-qPCR to analyze histone H4 expression and validate our observations in independent experiments using ZIKV. Histone H4 expression was significantly downregulated by ZIKV infection in the carcass of infected mosquitoes in a time-dependent manner (**Figure 5d**). In contrast, the presence of HTV and PCLV in mosquitoes prevented the downregulation of histone H4 following ZIKV infection (**Figure 5d**). In addition, levels of histone H4 were significantly higher in the presence of HTV and PCLV at every time point tested when compared to control mosquitoes (**Figure 5d**). We did not observe the same effect on the expression of histone H4 in the midgut, where HTV and PCLV did not affect susceptibility to ZIKV (**Supplementary Figure 9a**). We also observed that differential expression of histone H4 in the presence or absence of HTV and PCLV was only observed in DENV-infected mosquitoes (**Figure 5e**). These results suggest that HTV and PCLV prevent downregulation of histone H4 that is induced by infection with DENV and ZIKV. Accordingly, artificial infection of laboratory mosquitoes with HTV and PCLV alone did not significantly affect histone H4 expression levels (**Supplementary Figure 9b**).

**Fig. 5.**
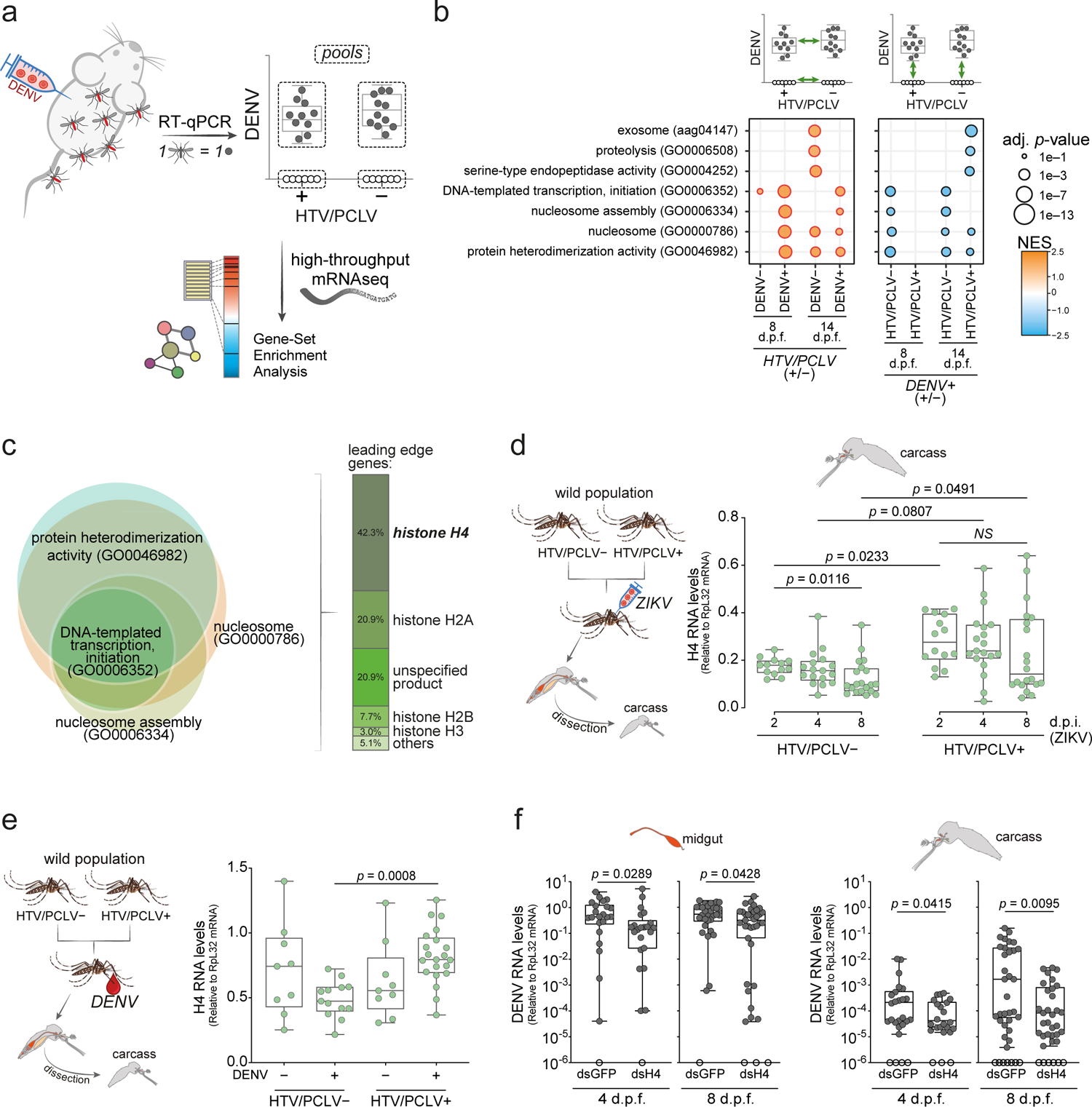
Histone H4 is a proviral host factor that is regulated by HTV and PCLV during DENV infection. **a**, Overall strategy to identify biological pathways associated with HTV and PCLV interaction with DENV. HTV/PCLV infected and virus free wild mosquito populations were exposed to DENV-infected mice for a blood meal. The transcriptome of DENV infected and non-infected mosquitoes from HTV/PCLV and virus-free groups were analyzed separately at 8 and 14 days post feeding. Gene Set Enrichment Analysis (GSEA) was performed for each comparison. **b**, Biological processes significantly enriched (adjusted *p*-value < 0.01) in the comparisons of DENV-infected versus non-infected mosquitoes and PCLV/HTV infected versus virus-free controls are shown. **c**, Overlap of leading-edge genes belonging to the 4 biological processes enriched in at least 6 out of 8 comparisons. Size of each circle represents the number of leading-edge genes. Histogram shows that histone genes represent the majority in the leading edge of significantly enriched processes. **d**, Histone H4 levels in the carcass of mosquitoes at 2, 4, and 8 days post injection of ZIKV in wild mosquitoes carrying HTV/PCLV or virus free controls. **e**, Differential expression of histone H4 levels in the carcass of mosquitoes in the presence of HTV and PCLV is only observed in DENV infected individuals. Carcass of mosquitoes exposed to oral infection with DENV analyzed at 8 days post feeding. **f**, Silencing of histone H4 mRNA decreases DENV replication in adult mosquitoes. DENV infection was analyzed at 4 and 8 days post blood feeding in the midgut and carcass of silenced (dsH4) and control (dsGFP) mosquitoes. *d.p.f.* – days post feeding. *d.p.i.* – days post infection, *NS* – non-significant. Boxplots represent minimum and maximum, median, and interquartile ranges of H4 mRNA levels (**d-e**) or detected viral load (**f**) for each group. Statistical significance was determined by ANOVA with Tukey’s correction for multiple comparisons (**d,e**) or by two-tailed Mann-Whitney U-test (**f**). Each dot represents an individual sample.

Overall, the above results raise the question of the role of histone H4 in DENV and ZIKV infection. In order to verify the importance of histone H4, we used dsRNA mediated gene silencing to target histone H4 in adult mosquitoes prior to infection (**Supplementary Figure 10**). Mosquitoes where histone H4 was silenced prior to infection showed reduced DENV levels at 4 and 8 days post infection (**Figure 5f**). Interestingly, silencing of histone H4 inhibited DENV replication in both midgut and carcass suggesting this is an important host factor for DENV replication regardless of the tissue (**Figure 5f**). However, since HTV and PCLV only affect histone H4 expression in the carcass, the effect of these ISVs over DENV and ZIKV replication is not observed in the midgut. Together, these results indicate that two ISVS prevalent in wild mosquitoes, HTV and PCLV, favor systemic dissemination of DENV and ZIKV by preventing the downregulation of histone H4, a novel proviral host factor.

## Discussion

Here we provide a global overview of viruses circulating in urban *A. aegypti* and *A. albopictus* mosquitoes that are responsible for the transmission of the most important human arboviruses such as DENV, ZIKV and CHIKV (*5*). Although metagenomics has been repeatedly applied to mosquitoes over the last few years (*8–13*), this is the first comparison of co-occurring urban *A. aegypti* and *A. albopictus* mosquitoes on a global scale. Our results identified a large diversity of ISVs circulating in *Aedes* mosquitoes, including 5 novel viral species. Despite the scale of our study, we did not find arboviruses in any of the *A. aegypti* and *A. albopictus* samples we analyzed. Since arboviruses generate as much of an RNAi response as ISVs (*9, 31–34*), it is unlikely that this is caused by a bias in detection in our small RNA-based strategy. Rather, our results confirm that arbovirus infections in wild mosquitoes are rare while ISVs are abundant and highly prevalent (*9, 10, 12, 13, 35, 37*).

The direct comparison of the virome in the two most important urban vectors for arboviruses, *A. aegypti* vs *A. albopictus*, suggests that they are very different in their ability to carry viruses. On a global scale and in each location, *A. aegypti* had higher prevalence of viruses and a more diverse virome than *A. albopictus*. While the virome of *A. aegypti* was often composed of 5-6 viruses in a single location, *A. albopictus* only carried a single virus at relatively low viral loads. This poor diversity in *A. albopictus* was observed in several independent samples, which suggests it is not an artefact. It is noteworthy that *A. albopictus* naturally carries the endosymbiont bacteria *Wolbachia* that is known to provide resistance to viruses in insects (*38–40*). Alternatively, the domestication and spread of *A. aegypti* happened a few centuries before that of *A. albopictus*, which could have allowed further diversification of viruses for the former species (*41*). Regarding the more diverse virome of *A. aegypti*, a total of 10 viral species were identified worldwide although there were local variations. Overall, the collection of viruses in mosquitoes was more similar among populations from geographically closer locations. Notably, the virome of African populations of *A. aegypti* was the most divergent from the rest. Even the most common *A. aegypti* viruses, PCLV and HTV, were found at lower abundance in Senegal and absent in Gabon mosquito populations even though they were highly prevalent in every other location. In the case of Gabon, this may be because mosquitoes belong to the *A. aegypti formosus* subspecies compared to the rest of the world where the *A. aegypti aegypti* subspecies dominates (*42*). Interestingly, the *A. aegypti formosus* subspecies is more resistant to arboviruses such as DENV and ZIKV (*43*), suggesting that the susceptibility to arboviruses and ISVs may be determined by similar genetic mechanisms. However, our results also establish that some ISVs render *A. aegypti* more susceptible to arboviruses, which could also explain the connection between the susceptibility to ISVs and arboviruses.

Our work provides strong field and laboratory evidence for the role of ISVs in shaping the transmission of arboviruses. Although many studies have reported interactions between ISVs and arboviruses, most were performed in cell lines (*21, 24–26*). Interestingly, in cell lines, PCLV either inhibited or did not affect the replication of ZIKV in cell lines (*20, 22*). Few studies in adult mosquitoes have focused on insect-specific flaviviruses such as Nhumirim virus (NHUV) and Cell fusing agent virus (CFAV) that were shown to interfere with replication of arboviruses of the same family such as West Nile virus, DENV and ZIKV (*17, 19, 23*). This is in stark contrast to our results, which show that the presence of PCLV favors arbovirus infections in adult mosquitoes. However, since we did not analyze PCLV alone in adult mosquitoes, we cannot rule out that HTV has a predominant effect. Indeed, our data from wild mosquitoes does suggest that HTV has the strongest association with DENV infection. These observations reinforce the importance of associating field studies with laboratory experiments in adult animals to be able to identify interactions between ISVs and arboviruses. Significant interactions might only be discovered by large field studies that are often expensive and do not usually analyze the circulation of ISVs. It is interesting to point out that one of our locations, the port city of Santos in the southeast of Brazil, showed co-circulation of PCLV, HTV and CFAV, which can affect the transmission of arboviruses in opposite directions. A large study in this location could provide insights on the interaction of multiple ISVs and arbovirus transmission in a city where DENV, ZIKV and CHIKV circulate. Overall, data on the virome of *A. aegypti* could help explain local variations in vector competence.

Our results that HTV and PCLV affect the susceptibility of *A. aegypti* mosquitoes to DENV and ZIKV led us to uncover the role of histone H4 as a novel proviral host factor. We observed that histone H4 was downregulated in response to DENV or ZIKV infection in mosquitoes but this is prevented by HTV and PCLV. Regarding how histone H4 functions as a proviral factor, it is noteworthy that the C protein from Flaviviruses is capable of interfering with nucleosome assembly (*44*). Recent work has suggested that the C protein of Yellow fever virus and possibly other flaviviruses mimics the tail of histone H4 and regulates gene expression to favor infection (*45*). Thus, downregulation of histone H4 may be part of a coordinated host response to limit the ability of flaviviruses to control gene expression, and this is counteracted by HTV and PCLV. These results open new perspectives for the field and raise fresh questions, about how HTV and PCLV regulate histone H4 during DENV and ZIKV infection and whether this mechanism affects arboviruses from other families such as CHIKV. Importantly detailed mechanistic understanding about how ISVs affect vector competence could point to alternative strategies of controlling the transmission of arboviruses.

## Acknowledgments

We thank all members of the Zikalliance consortium, especially Anna-Bella Failloux and Alain Kohl, that helped establish a network of collaborators and contributed with fruitful discussions. We also thank all members of the Marques laboratory and the M3i unit - Insect Models of Innate Immunity, especially Stephanie Blandin, for suggestions and discussions.

## Author contributions

Conceptualization: RPO, ERGRA, JLI, JTM Funding acquisition: ERGRA, MLN, JLI, JTM

Methodology: RPO, YMHT, ERGRA, JPPA, JNA, LS, JTM

Investigation: RPO, YMHT, ERGRA, JPPA, JNA, IJSF, FVF, ATSS, KPRS, APPV, LS, JTM

Validation: RPO, YMHT, ERGRA, JPPA, JNA, IJSF, FVF

Data curation: RPO, YMHT, ERGRA, JPPA, JNA, LS, JTM Formal analysis: RPO, YMHT, ERGRA, JPPA, JNA, LS, JTM Visualization: RPO, YMHT, ERGRA, JPPA, JNA, JTM

Resources: CHT, MD, AG, CP, JON, TMV, CJMK, MAW, ALCC, MTP, MCPP, MLN, VAS, RNM, MAZB, BPD, EM

Project administration: RPO, JLI, JTM Supervision: JLI, JTM

Writing – original draft: RPO, YMHT, ERGRA, JLI, JTM

Writing – review & editing: RPO, YMHT, ERGRA, JPPA, CP, TMV, CJMK, MLN, VAS, BPD, LS, JLI, JTM

## Data availability

Small RNA libraries and transcriptome libraries from this study have been deposited on the Sequence Read Archive (SRA) at NCBI under the project accession PRJNA722589. Other publicly available RNA-seq data sets were obtained from SRA. Accession numbers for small RNA libraries are provided in **Supplementary Table 1**.

## Supplementary Material

### Materials and Methods

#### Ethics statement

All procedures involving vertebrate animals were approved by the ethical review committee of the Universidade Federal de Minas Gerais (CEUA 337/2016 to J.T.M.).

#### Sample collection in the field

Locations of mosquito collections are described in the **Supplementary Table 1**. Field traps were used to collect adult mosquitoes that were further identified using morphological characteristics. Whole mosquitoes were grinded in TRIzol (Invitrogen) and kept refrigerated prior to RNA extraction.

#### RNA extraction

RNA was isolated using TRIzol reagent (Invitrogen) following the manufacturer protocol with minor modifications. Briefly, individual mosquito samples or tissues were collected in 1,5 mL tubes, 3-5 glass beads (1 mm diameter) and ice-cold TRIzol were added before being homogenized in a Mini-BeadBeater-16 (Biospec^©^) for 90 seconds. Glycogen (Ambion) was added (10µg per sample) to facilitate pellet visualization upon RNA precipitation. RNAs were resuspended in RNAse-free water (Ambion) and stored at −80°C.

### Small RNA library construction

Different strategies for library construction were implemented and are indicated in **Supplementary Table 1**. The strategy was determined according to RNA quality evaluated by the 2100 Bioanalyzer system (Agilent). Libraries were built using total RNA or size selected small RNAs (18-30 nt), depending on quality and yield of the sample. In the case of low RNA yield, especially when the source was a single mosquito, total RNA was directly used as input for library preparation. For samples with more than 20 ug of RNA available, small RNAs were selected by size (18–30 nt) on a denaturing PAGE. For samples with more than 20 ug of total RNA that displayed a degradation profile (*i.e.*, lack of sharp ribosomal RNA peaks), total RNA was subjected to oxidation using sodium periodate (*46, 47*), prior to size selection. Oxidized and non-oxidized size selected RNAs (18-30 nt) were used as input for library construction. In all cases, libraries were prepared utilizing the TruSeq^®^ Small RNA Library Prep Kit (Illumina^®^) or NEBNext^®^ Multiplex Small RNA Library Prep Set for Illumina^®^ (New England BioLabs inc.) following protocols recommended by the manufacturers. Libraries were pooled and sequenced at the GenomEast sequencing platform at the Institut de Génétique et de Biologie Moléculaire et Cellulaire in Strasbourg, France.

### Small RNA-based metagenomics for virus identification

After sequencing, raw sequenced reads from small RNA libraries were submitted to adaptor trimming using cutadapt (*48*) v1.12, discarding sequences with low Phred quality (< 20), ambiguous nucleotides and/or with length shorter than 15 nt. Remaining sequences were mapped to reference sequences of *A. aegypti* (AaeL5) (*49*) or *A. albopictus* (*50*) using Bowtie (*51*) v1.1.2 allowing no mismatches. Size profiles of small RNAs matching reference sequences and 5’ nt frequency were calculated using in-house Perl v5.16.3, BioPerl library v1.6.924 and R v3.3.1 scripts. Plots were made in R using ggplot2 v2.2.0 package. Sequences that did not present similarities with bacteria or the host genomes were used for contig assembly and subsequent analyses. Assembly was performed essentially as previously described (*9*) with the following changes: (1) We replaced Velvet (*52*) assembler by SPAdes (*53*) on the second round of contig assembly. (2) Assembled contigs ranging from 50 to 199 nt were characterized solely based on sequence similarity search against Viral RefSeq Database (*54*). (3) Contigs greater than 200nt were characterized based on sequence similarity against the NCBI NT and NR databases and submitted to pattern-based strategies. (4) For manual curation of putative viral contigs, top 5 BLAST (*55*) hits were analyzed to rule out similarity to other organisms, ORF organization and small RNA size profile (distribution and coverage) were analyzed to differentiate between viruses. Contigs containing truncated ORFs and small RNA profiles without the presence of symmetric small RNA peaks at 21 nt were considered to be EVEs as described (*30*). (5) Manually curated viral contigs were grouped using CD-HIT (*56*) requiring 90% of coverage with 90% of identity to remove redundancy. Representative contigs were used for co-occurrence analysis based on small RNA abundance on each of the small RNA libraries available. Contigs grouped into a single cluster (Hierarchical clustering based on Pearson correlation) were then used as trusted on SPAdes for a re-assembly step using all the libraries in which that viral sequence was found. For more details on manual curation steps, see Figure 2A.

### Phylogenetic analyses

Assembled viral contigs were submitted to analysis of conserved domains to identify RdRp-related regions using NCBI Conserved Domain Search (https://www.ncbi.nlm.nih.gov/Structure/cdd/wrpsb.cgi). For each putative virus, the largest RdRp segment was used to identify virus relatives at NCBI sequence databases (NT and NR) using sequence-similarity searches through BLAST tool. Multiple alignments were performed using the MAFFT online tool (*57*) available at https://www.ebi.ac.uk/Tools/msa/mafft/. For putative new viruses identified at protein level (*Narnaviridae, Partitiviridae, Rhabdoviridae, Totivirdae* and *Virgaviridae*), amino acid sequences were selected, and phylogenetic analyses were carried out on CIPRES Portal version 3.3 (https://www.phylo.org/portal2) (*58*). The best-fit model of protein evolution was selected in ModelTest-NG (*59*) for each viral species, using Maximum Likelihood (ML) method. For the virus from *Totiviridae* family, an additional strategy was also applied using nucleotide sequences where the best-fit model was defined using MEGA-X tool (*60*), and the tree was constructed using ML method. For all phylogenetic trees, clade robustness was assessed using bootstrap method (1000 pseudoreplicates) and trees were edited using iTOL version 5.7 (*61*) (https://itol.embl.de/).

### Mosquitoes

Wild *A. aegypti* mosquitoes used in experiments described here were F2 to F5 generations derived from eggs collected in the Rio de Janeiro city (Urca neighborhood) in Brazil and were kindly provided by Dr. Rafael M. de Freitas from Fiocruz-RJ and Dr. Luciano A. Moreira from Fiocruz-MG. The laboratory Red-eye strain (*62*) was kindly provided by Prof. Pedro C. Oliveira from the Universidade Federal do Rio de Janeiro - UFRJ. *A. aegypti* mosquitoes were maintained in a climatic chamber at 28°C and 70-80% relative humidity, in a 14:10 hour light:dark photoperiod, and 10% sucrose solution *ad libitum*. Mosquito cages were composed of individuals that emerged within an interval of 48 hours.

### Generation of HTV+/PCLV+ and HTV–/PCLV– mosquito lines

Mosquito lines persistently infected with PCLV and HTV or non-infected counterparts were generated from F2 generations of wild mosquitoes. Three days after a blood meal, F2 mated females were individually isolated in tubes containing a filter paper and 0.5 cm of water and were allowed to lay eggs for 24 hours. Individual females were collected, and the total RNA was isolated using TRIzol (Invitrogen) following the standard protocol. Detection of HTV and PCLV was performed by RT-qPCR using the primers described in **Supplementary Table 5**. Eggs corresponding to 5 female mosquitoes infected with HTV, PCLV or both viruses were pooled prior to hatching. Pools of eggs from 5 females negative for both viruses were pooled similarly to create virus free lines. Subsequent detection of HTV and PCLV was performed to confirm correct identification of lines. Lines generated from females carrying a single virus (HTV or PCLV) were tested and found to carry both viruses. Therefore, only virus free and double infected lines were expanded for two more generations for experiments described in this work.

### Artificial infection of naïve laboratory *A. aegypti* mosquitoes with HTV and PCLV

Extracts of naturally infected *A. aegypti* mosquitoes were used as source for HTV and PCLV since we were not able to produce these viruses in cell culture. Viral stocks were produced from pools of 15 *A. aegypti* naturally infected with HTV and PCLV or non-infected mosquitoes (virus-free controls), that were grinded using pestles in 1200 µL of L-15 Leibovitz medium (Gibco) supplemented with 10% fetal bovine serum (FBS). Samples were centrifuged at 3000 × g for 15 minutes at 4°C for clarification. Supernatants were collected and passed through a 0.22 µm filter, aliquoted, and stored at −80°C prior use. Infection with HTV and PCLV or mock control was performed by microinjecting 69 nL of the extract into naïve laboratory mosquitoes (*A. aegypti* RedEye strain) using a Nanoject II microinjector (Drummond).

### Infection of Vero cells with HTV and PCLV

Filtered *A. aegypti* extracts (500 µL) containing HTV and PCLV (obtained as described above) were transferred into T-25 flasks containing 90% confluent Vero cells in non-supplemented DMEM medium. After one hour of viral adsorption, 4.5 mL of DMEM medium supplemented with penicillin/streptomycin and 10% of FBS were added to cells, that were incubated at μL aliquots of the supernatant were collected during each passage on days 1, 3 and 5 after exposure to HTV and PCLV. Virus in the supernatant was assessed by RT-qPCR. Vesicular stomatitis virus (VSV) was added as a spike immediately before RNA extraction to be used as an internal control.

### DENV and ZIKV propagation

Viral isolates of DENV1 (isolate MV09) and ZIKV (PE243/2015)(*63*) were propagated in C6/36 *A. albopictus* cells or Vero cells respectively. For DENV1 propagation, C6/36 cells were maintained on L15 medium supplemented with 5% FBS (fetal bovine serum) and 1x Antibiotic-Antimycotic (Gibco) as described(*35*). Cells were seeded to 70% confluence and infected at a multiplicity of infection (MOI) of 0.01 and maintained for 6 to 9 days at 28°C. For ZIKV propagation, a similar procedure was followed using Vero cells that were maintained in DMEM medium supplemented with 10% FBS (fetal bovine serum) and 1x Antibiotic-Antimycotic (Gibco). Vero cells were seeded to a confluence of 70-80% infected with ZIKV at a MOI of 0.01 and maintained for 6 days in culture. For both viruses the supernatant was collected and clarified by centrifugation to generate virus stocks that were kept at −80°C prior to use. Mock-infected supernatants used as controls were prepared under same procedure without virus infection. Titration of DENV1 was performed in BHK-21 cells while ZIKV was titrated in Vero cells, both using the plaque assay method to determine viral titer.

### DENV and ZIKV infection in mosquitoes

Natural infection in mosquitoes was performed using mice deficient for interferon-I and interferon-II receptors as described (*34*). Briefly, infection in AG129 mice was established by intraperitoneal injection of approximately 10^6^ PFU of DENV1 or 10^6^ PFU of ZIKV. Infected mice were anaesthetized 3 days post infection (peak of viraemia) using ketamine/xylazine (80/8 mg per kg) and placed on top of the netting-covered containers with adult mosquito females. Mosquitoes were allowed to feed on mice for 1h, with mice rotation intervals of 10 min between containers. For infections by membrane feeding, 5-6 day old adult females were starved for 24h and fed with a mixture of blood and virus supernatant containing 10^7^ PFU/mL of DENV serotype 1 utilizing a glass artificial feeding system covered with pig intestine membrane as previously described (*34*). After blood feeding, fully engorged females were selected and kept in standard rearing conditions until collection at different time points. Mosquitoes infected by injection were anesthetized with cold at 4°C and kept on ice during the whole procedure. Virus stocks were diluted with L15 medium (Gibco) and injections were carried out using the microinjector Nanoject II (Drummond) with a volume of 69 nL. Each individual mosquito was injected with 16 PFU of DENV or ZIKV. Mosquitoes were harvested at different days post injection for dissection and RNA extraction. Tissues (midguts or ovaries) were dissected in ice-cold 1x phosphate buffered saline (PBS) containing 0.01% Triton X-100 (Sigma). Remnants of mosquito tissues were considered as carcass, as illustrated in figure schemes. Tissues or mosquitoes were ground in TRIzol (Invitrogen) using glass beads as previously described (*34*). Total RNA was extracted from individual mosquitoes or individual tissues according to manufacturer’s protocol.

### RT-qPCR and RT-PCR

Total RNA extracted from individual mosquitoes or individual tissues were reverse transcribed using M-MLV reverse transcriptase using random primers for initiation. Negative controls were prepared following the same protocol without adding the reverse transcriptase. cDNA was subjected to quantitative polymerase chain reaction (qPCR) utilizing the kit Power SYBR^®^ Green Master Mix (Applied Biosystems) following manufacturer instructions. Results were expressed using the 2-ΔCt method relative to the endogenous control *rpL32*. Primer sequences are listed in **Supplementary Table 5**.

### Poly-A selection, RNA library construction, and transcriptomic analysis

RNA samples from individual mosquitoes were pooled and RNA quality was verified using the 2100 Bioanalyzer system (Agilent). mRNA libraries were constructed using the kits NEBNext^®^ Poly(A) mRNA Magnetic Isolation Module and NEBNext^®^ Ultra^™^ II Directional RNA Library Prep Kit for Illumina^®^ (New England BioLabs inc.) following the manufacturer protocol. Libraries were pooled and sequenced at the GenomEast sequencing platform at the Institut de Génétique et de Biologie Moléculaire et Cellulaire in Strasbourg, France. Sequenced reads with an average quality score above phred 25 had adaptors removed using Trimmomatic v0.39 and were further mapped to the decoyed transcriptome of *A. aegypti* (Vectorbase release 48) using Salmon v1.3.0 (*64, 65*). Quasi-mapping quantifications were imported into R v3.6.3 and data normalization was performed using the packages EdgeR v3.28.1 and TMM (*66, 67*). Ranked lists of gene expression for each comparison was used as input for Gene Set Enrichment Analysis (GSEA) (*36*) using the R package fgsea v1.12.0 (*68*) and in-house developed gene-sets comprising Gene Ontology annotation, pathways, and genes of interest. Sets with adjusted *p*-value < 0.1 were considered in our analysis.

## Statistical analyses

Evaluation of statistical significance was performed using R-cran v3.3.1 software unless stated otherwise. Viral loads of RT-qPCR positive-only individual mosquito/tissues were log transformed and subjected to Mann-Whitney *U* test. Prevalence of infection was evaluated by Fisher’s exact test. Presence and absence of DENV in mosquitoes was modelled with univariate and multivariate zero-inflated binomial model (ZIB) (*69, 70*), since 95% of the collections are zeroes. The covariate or covariates (in case of multivariate model) was/were the same for the logit and logistic parts of the model. In particular, we considered PCLV, HTV and their interaction. Model selection was carried out by AIC and BIC comparison (*71*). The Vuong test was conducted a priori to test if the zero inflated binomial model was statistically significant and better (in terms of AIC and BIC) than a non-zero inflated model. Data were analysed using R and the ‘pscl’ package v. 1.5.5 (*72*). Finally, we tested the presence of spatial autocorrelation in the two viruses – via variogram analyses (*35*) - but no significant autocorrelations have been found (results not reported).

## Supplementary Figures

**Supplementary figure 1.**
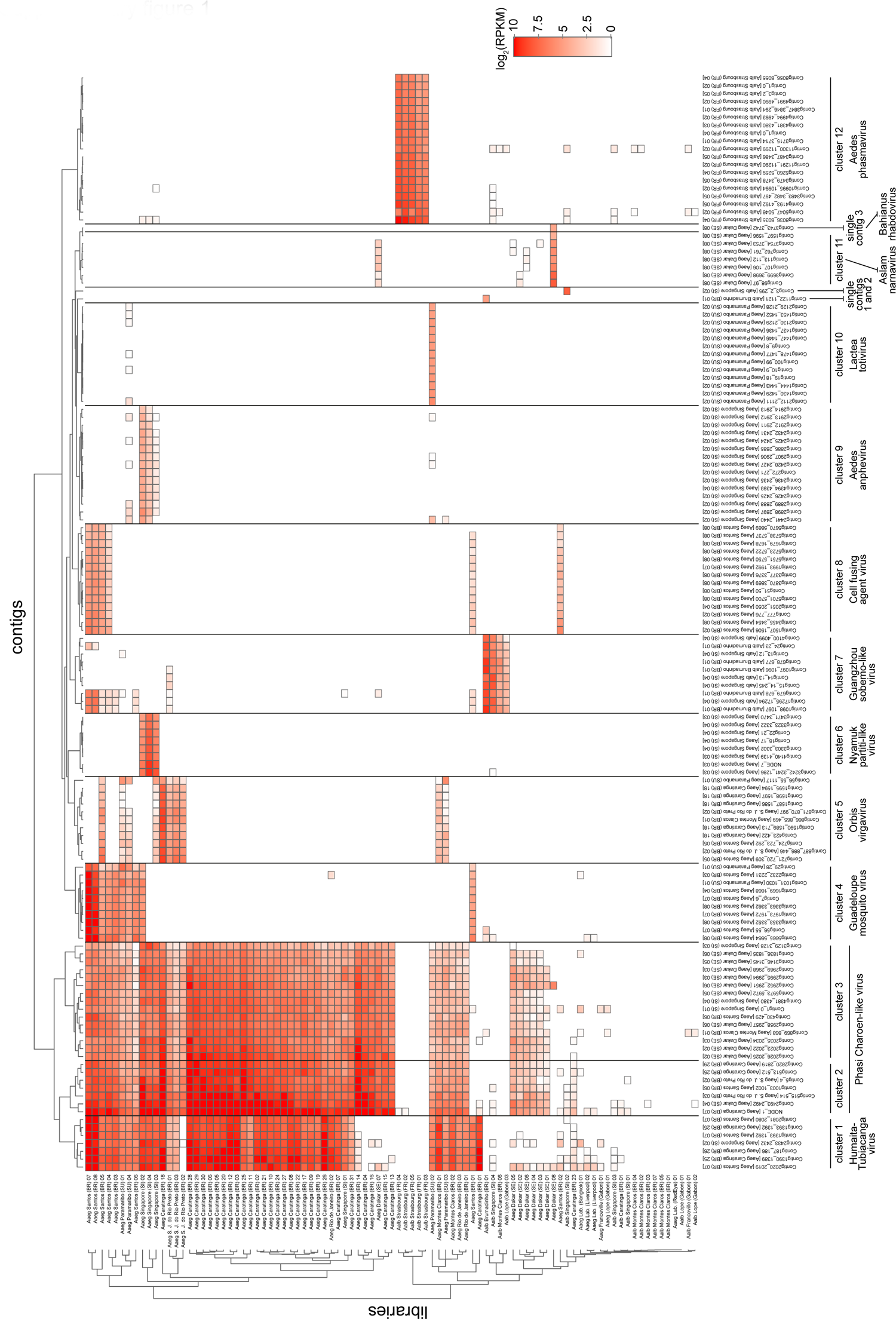
Co-occurrence of 139 viral contig clusters identified in *A. aegypti* and *A. albopictus* mosquitoes. Heatmap represents the small RNA abundance for each contig. White indicates absence of small RNAs mapping to that contig. Contig clusters were defined using the dendrogram above the heatmap. Clusters that had a RdRp sequence were classified as a putative virus, that was only considered present in a sample if >50% of contigs belonging to a cluster were represented.

**Supplementary figure 2.**
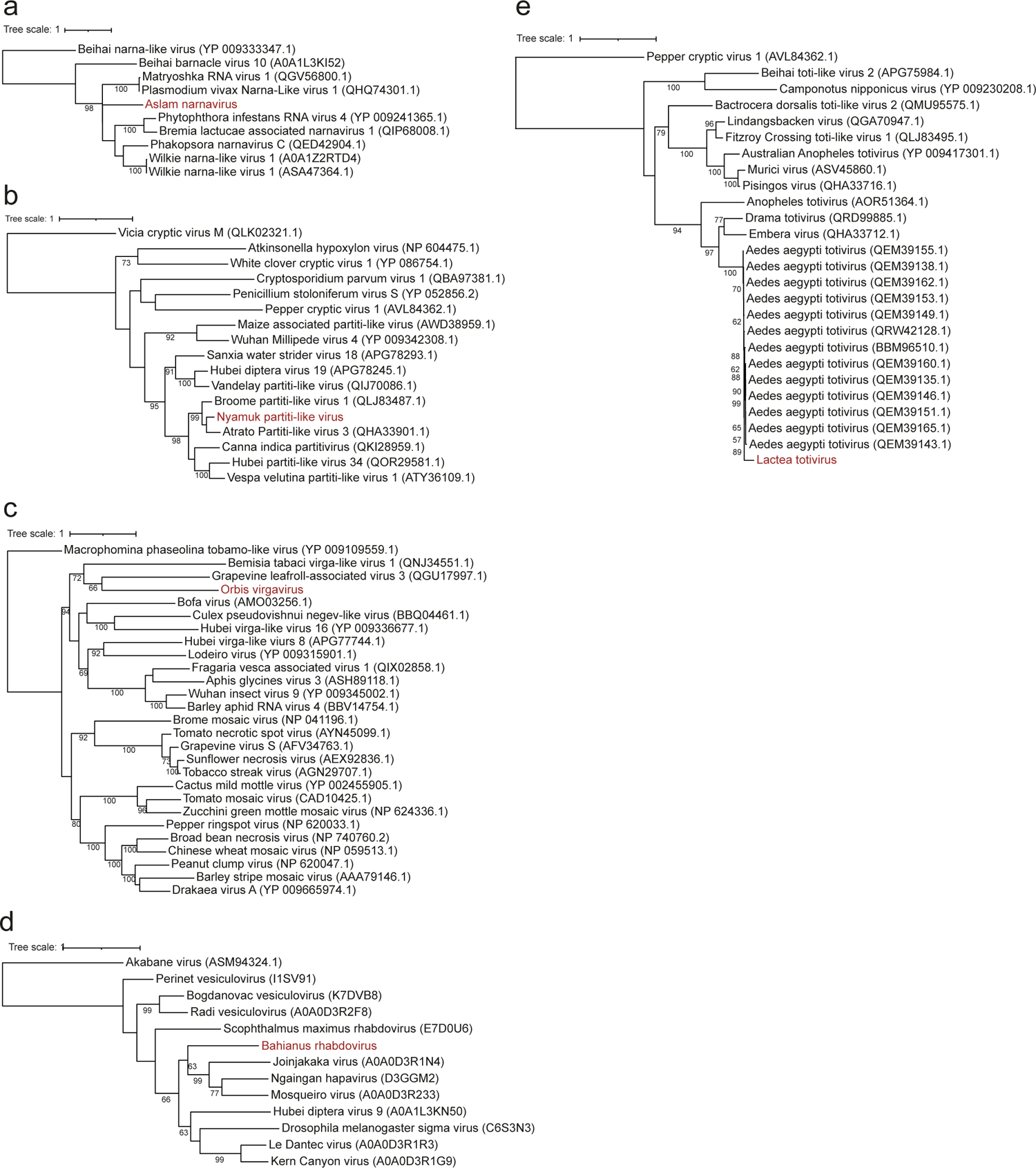
Phylogeny of viruses identified in *A. aegypti* mosquitoes. Phylogenetic trees were generated using the RdRp amino acid (aa) or nucleotide (nt) sequences and the substitution models as indicated: **a**, Aslam narnavirus (aa - LG + G); **b**, Nyamuk partiti-like virus (aa - BLOSUM 62); **c**, Orbis virgavirus (aa - BLOSUM62 + F); **d**, Bahianus rhabdovirus (aa - BLOSUM62); **e**, Lactea totivirus (nt - Tamura-Nei 93). Bootstrap confidence is shown close to each clade and values under 60% were omitted.

**Supplementary figure 3.**
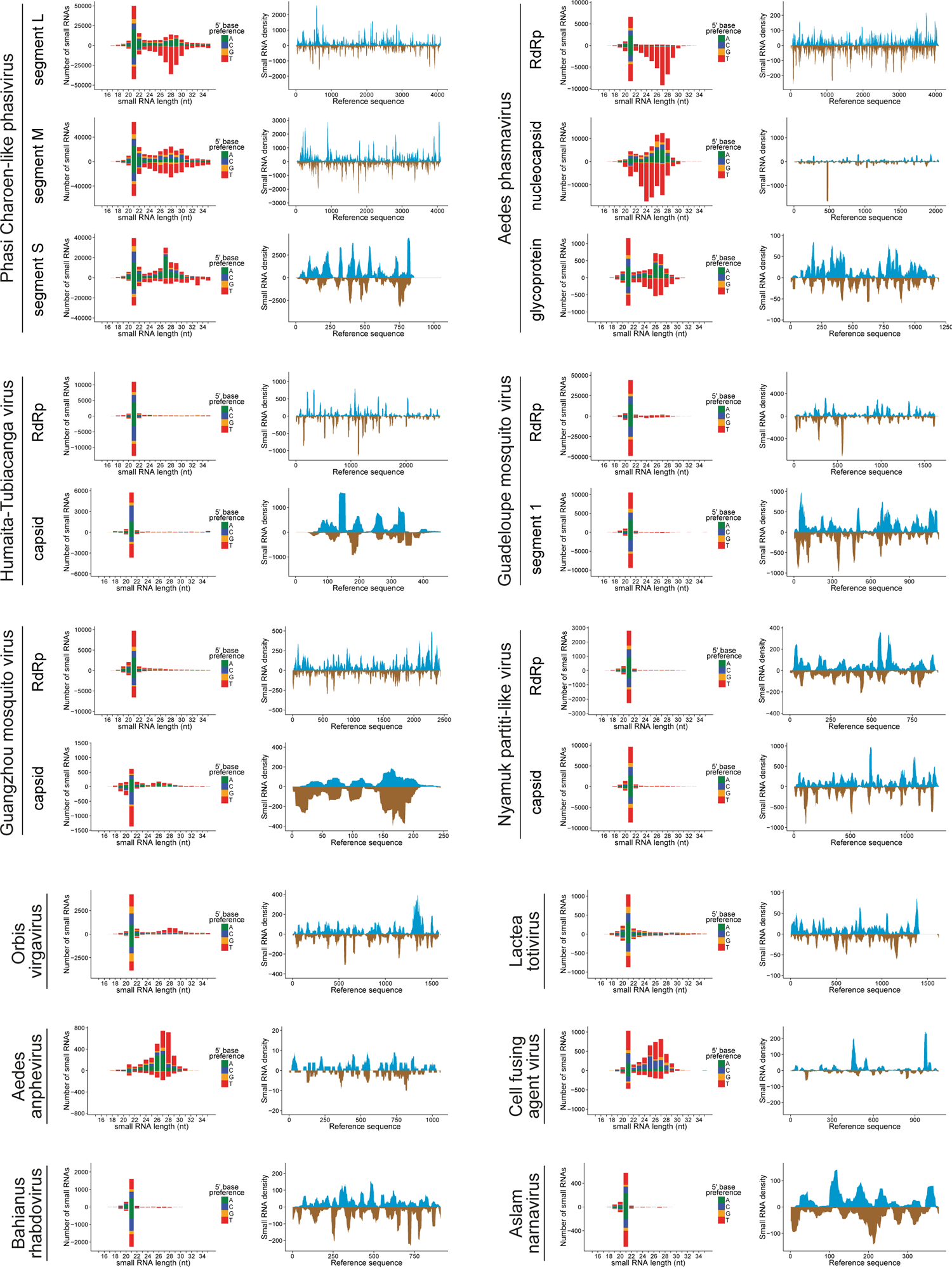
Virus-derived small RNA profiles in mosquitoes. Small RNA size distribution and 5’ base preference is shown on the left while the density of small RNAs (coverage) is shown on the right for representative contig(s) of each of the 12 viruses identified in this study.

**Supplementary figure 4.**
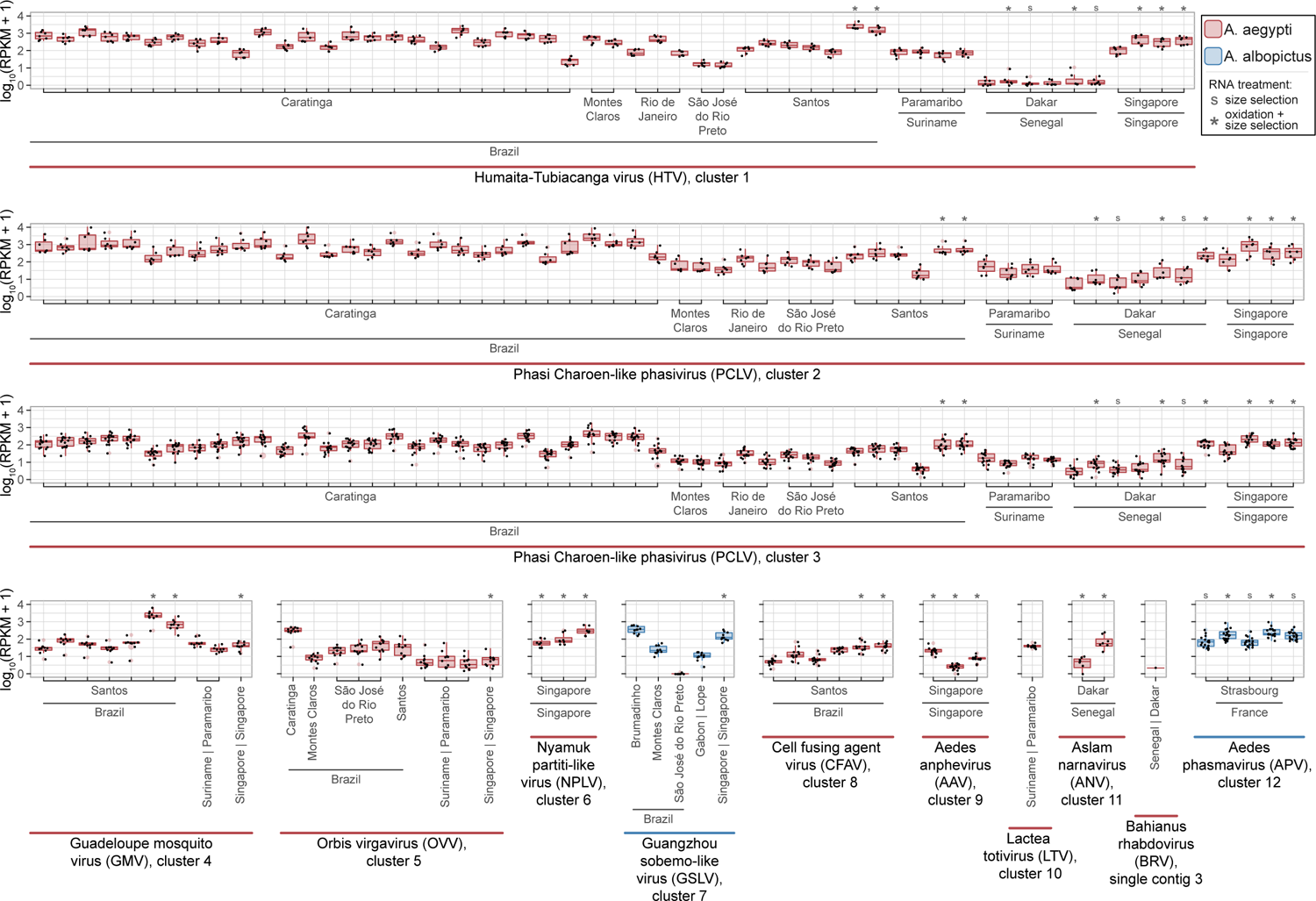
Burden of viruses in mosquitoes from different collection sites. Each dot represents the small RNA abundance in a contig, and boxplots represent contig clusters (see Supplementary Figure 1) at different locations with colors matching the mosquito species. Boxplots represent average, interquartile interval, and standard deviation of the mean for each location. RNAs that were size selected (18-30 nt) prior small RNA library construction are indicated with “S” and RNAs that were oxidized and size selected are indicated with “*”.

**Supplementary figure 5.**
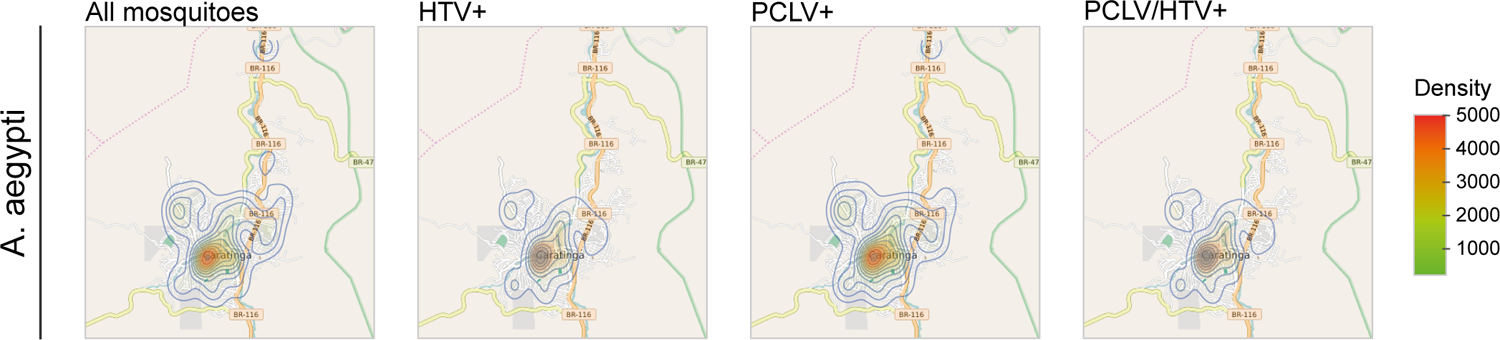
Geographic distribution of mosquitoes carrying HTV and PCLV in the city of Caratinga, Brazil. Maps show the density of adult *A. aegypti* mosquitoes captured in the city of Caratinga from July 2010 until August 2011 estimated from the number of mosquitoes captured in individual traps. All mosquitoes, HTV positive, PCLV positive and double positive individuals are shown. Viral detection was performed by RT-qPCR.

**Supplementary figure 6.**
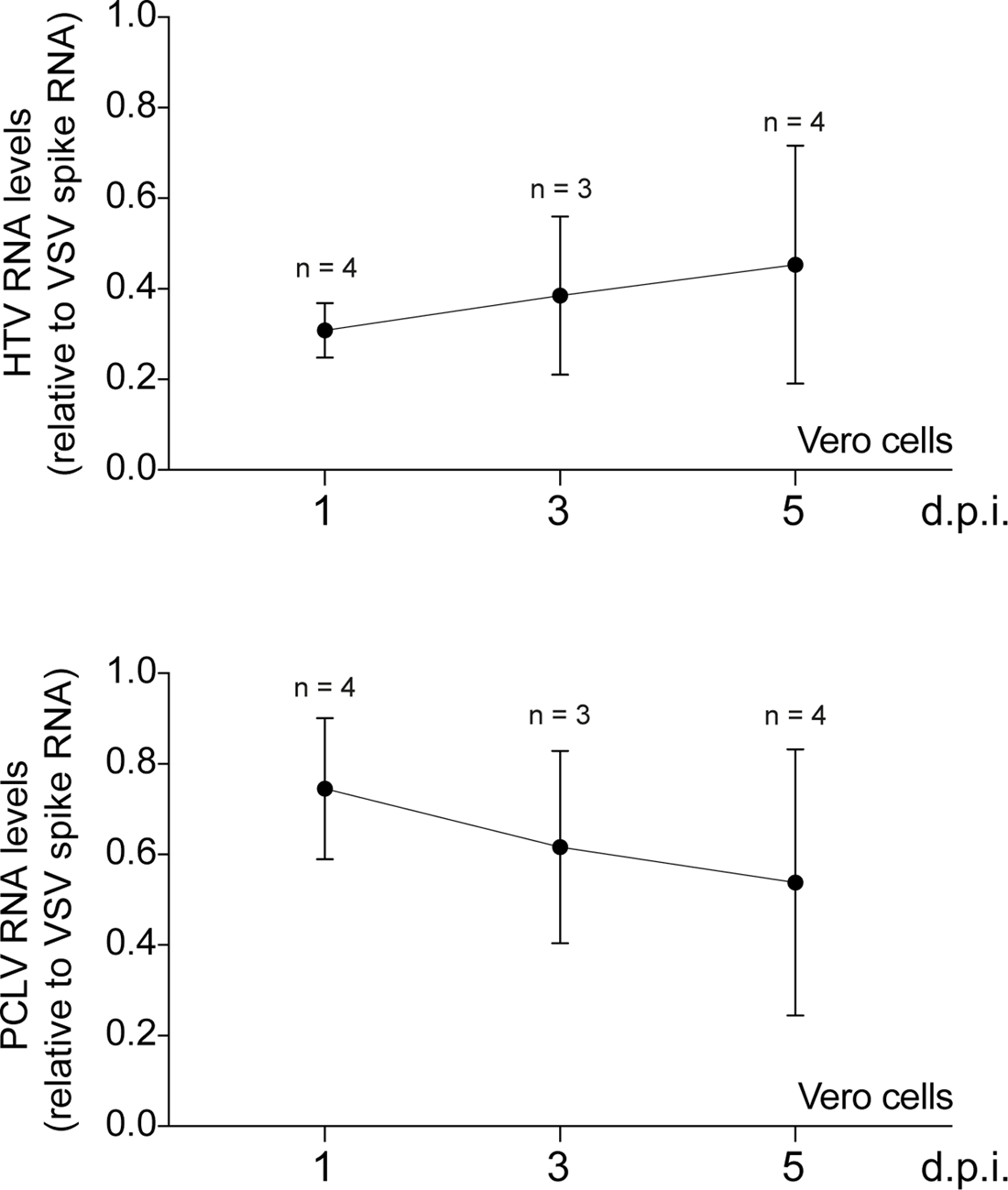
HTV and PCLV do not grow in mammalian cell culture. VERO cells were exposed to mosquito extracts containing HTV and PCLV, and supernatants were collected at 1, 3 and 5 days post exposure. A spike containing 10^5^ pfu of Vesicular stomatitis virus (VSV) was added prior RNA extraction and used to normalize the quantification of HTV and PCLV in the supernatant. No statistically significant difference was observed in HTV and PCLV levels at 1, 3 and 5 days post infection as determined by ANOVA with Tukey’s correction for multiple comparisons.

**Supplementary figure 7.**
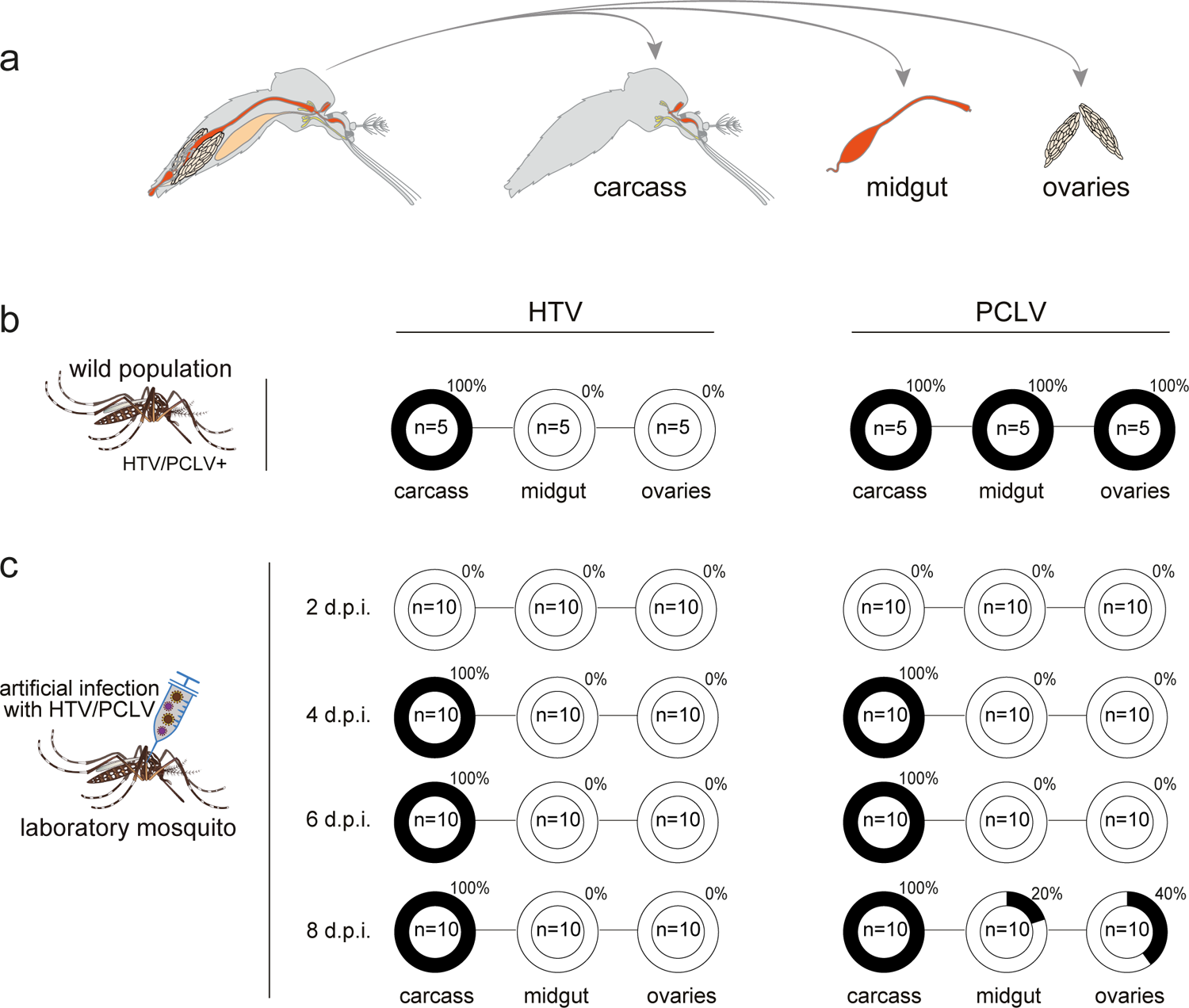
Tissue tropism of HTV and PCLV upon natural and artificial infections in *A. aegypti* mosquitoes. **a**, Scheme of mosquito dissection and tissues tested for virus infection by RT-qPCR. **b**, Prevalence of HTV and PCLV infection was assessed in a wild population of mosquitoes persistently infected with both viruses. **c**, Laboratory virus-free mosquitoes were injected with HTV and PCLV and the infection with both viruses was assessed at 2, 4, 6 and 8 days post injection (*d.p.i.*).

**Supplementary figure 8.**
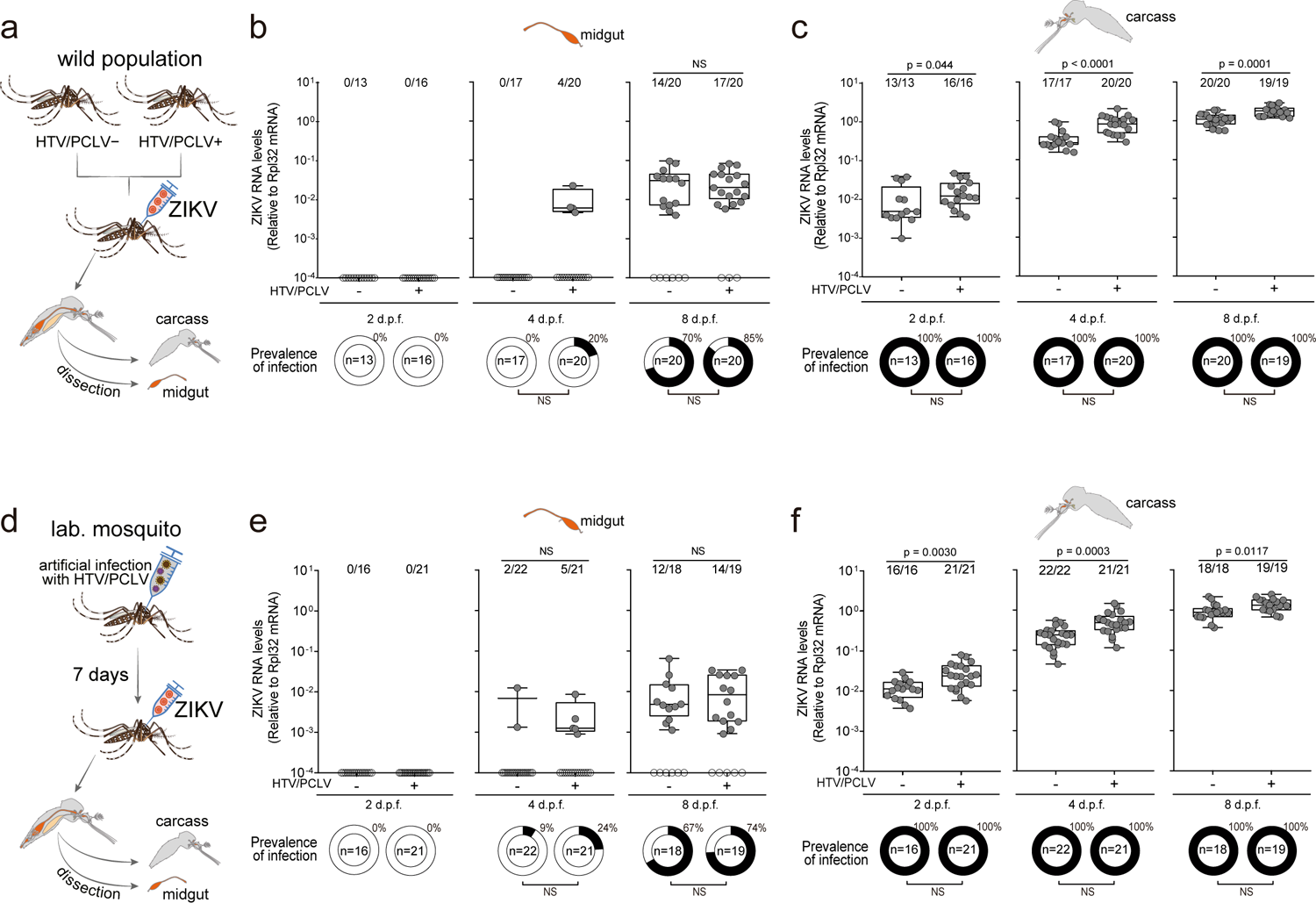
HTV and PCLV facilitate direct systemic ZIKV infection in mosquitoes. **a-c**, Strategy to evaluate the interference of HTV and PCLV for ZIKV infection and replication in natural populations of mosquitoes. (**a**) HTV/PCLV infected and virus-free wild mosquito populations were exposed to ZIKV by injection with the virus. Viral loads and prevalence of infection were measured in the (**b**) midgut and (**c**) carcass of mosquitoes at 4, 8 and 14 days post feeding. The prevalence of infection in each group is shown below the plots. **d-f**, Laboratory mosquitoes (**d**) were infected artificially with HTV and PCLV and 7 days later were injected with ZIKV. Viral loads and prevalence of infection were measured in the (**e**) midgut and (**f**) carcass of mosquitoes at 2, 4 and 8 days post injection. The prevalence of infection in each group is shown below the plots. *d.p.f.* – days post feeding, *NS* – non-significant. In ***b***, ***c***, ***e***, and ***f***, error bars represent median and interquartile range, and statistical significance was determined by two-tailed Mann–Whitney U-test. Numbers of infected midguts (***b***,***e***) or carcasses (***c***,***f***) over the total number tested are indicated above each column. Each dot represents an individual sample. Statistical significance of prevalence was determined by two-tailed Fisher’s exact test.

**Supplementary figure 9.**
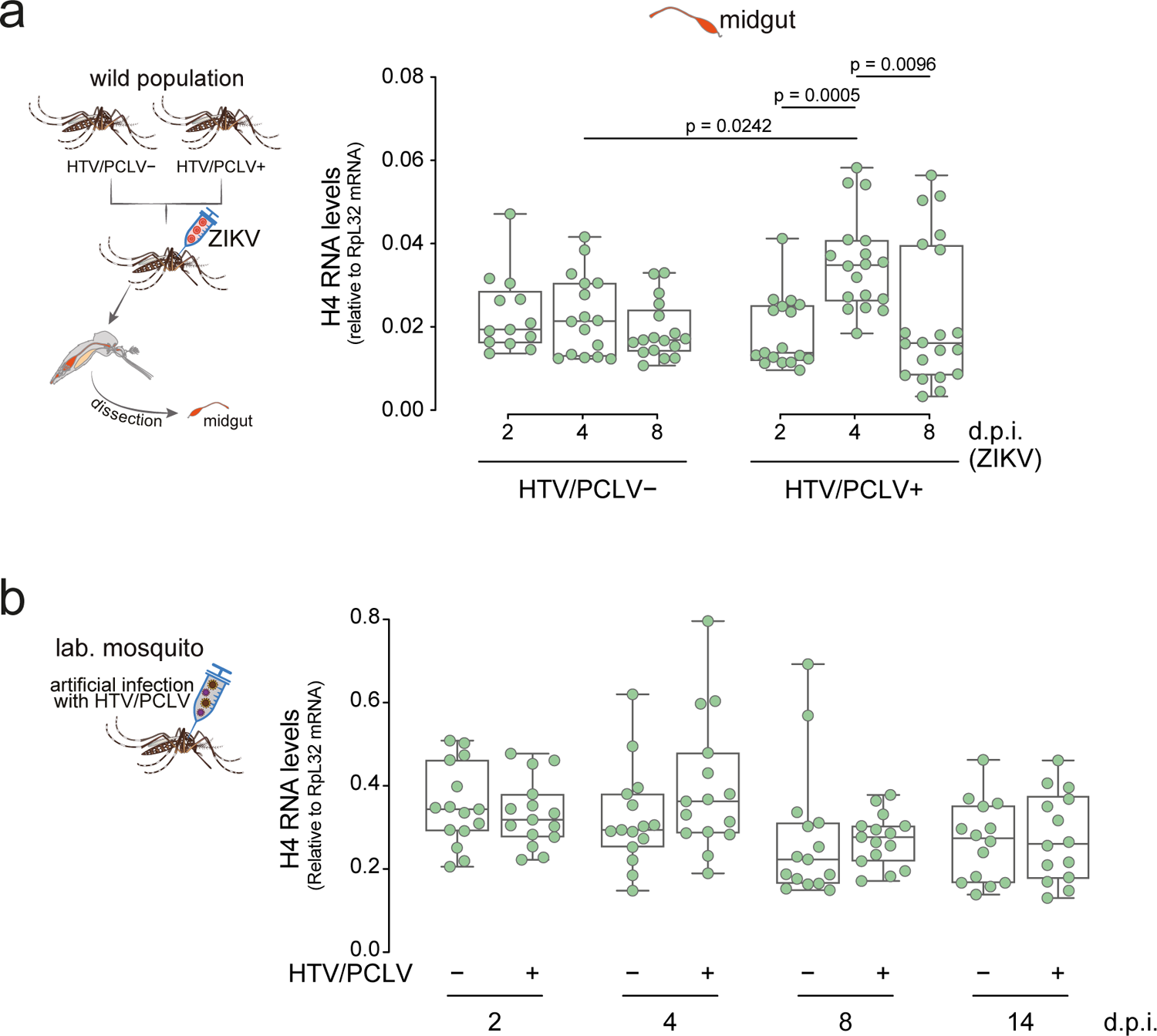
Histone H4 expression in wild mosquitoes carrying HTV and PCLV or artificially infected laboratory mosquitoes. **a**, Histone H4 levels in the midgut of mosquitoes at 2, 4, and 8 days post injection with ZIKV in wild mosquitoes carrying HTV/PCLV or virus free controls. *d.p.i.* – days post infection, *NS* – non-significant. Error bars represent mean and standard deviations of the mean, and statistical significance was determined by one-way ANOVA with Tukey’s correction for multiple comparisons. Each dot represents an individual sample. **b**, Artificial infection of laboratory mosquitoes by HTV and PCLV does not modulate levels of histone H4. Mosquitoes were infected with HTV and PCLV by injection and histone H4 levels were analyzed at different days. No comparison showed statistical significance between groups when using ANOVA with Tukey correction for multiple comparisons.

**Supplementary figure 10.**
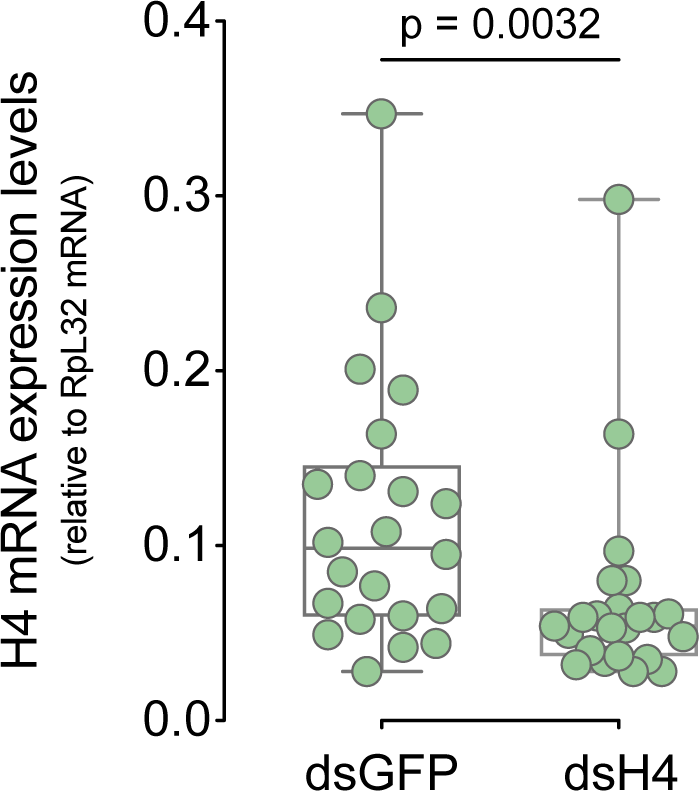
Silencing of histone H4 by RNA interference in adult mosquitoes. Histone H4 levels in mosquitos injected with dsRNA targeting GFP (dsGFP) as control or histone H4 (dsH4) at 4 days post feeding. Statistical significance was determined by two-tailed Mann-Whitney U-test.

## Supplementary Tables

**Supplementary table 1 – Overview of small RNA libraries**

**Metadata of small RNA libraries used in our study (SRA deposit ID, species, mosquito capture location, number of mosquitoes per pool, RNA treatment, and sequencing method).**

**Supplementary table 2 – Small RNA assembly metrics**

**Detailed information of assembled contigs with length >50nt and length >199 nt per small RNA library.**

**Supplementary table 3 – Overview of contigs with similarity to viral sequences deposited in GenBank**

**Detailed information of BLASTn or BLASTp hits from contigs matching viral sequences (length >199 nt) per small RNA library.**

**Supplementary table 4 – Overview of CDHit clusters**

**Number of contigs that compose each CDHit cluster.**

**Supplementary table 5 – List of oligonucleotides used in this study**

**Oligonucleotide sequences used for RT-qPCR.**

